# Drosophila Undigested Metabolite Profiling reveals age related loss of intestinal amino acid transport regulates longevity

**DOI:** 10.1101/2023.10.26.564159

**Authors:** Abigail A. Mornement, Rachael E. Dack, David P. Doupé, Rebecca I. Clark

## Abstract

Age-related intestinal decline is marked by altered epithelial architecture, loss of barrier function, elevated stress and immune signalling and changes to the intestinal microbiota. Despite this we do not yet know whether age-related intestinal decline impacts nutrient management, a key function of the intestinal epithelium.

In this study we have developed *Drosophila* Undigested Metabolite Profiling (D.U.M.P.) to assess the impact of intestinal ageing on nutrient absorption/excretion balance. We demonstrate that ageing results in a significant increase in amino acid load in the faecal matter that is largely driven by the microbiota and shortens lifespan. Increased amino acid load is associated with reduced expression of a subset of amino acid transporters. Knockdown of the amino acid transporter *slimfast* in the intestinal epithelium extends lifespan and confers improved microbial control in aged flies, suggesting reduced transporter expression is protective, preventing cellular uptake of excess amino acids.

We conclude that age-related changes to the microbiota are an important determinant of the local nutritional environment, with consequences for health. In addition, age-related decline of the intestinal epithelium may impact its capacity for nutrient absorption. These findings have significant implications for the rational design of anti-ageing nutritional therapies.

## Introduction

The ageing intestine has been the subject of intensive study in recent years with significant advances in our understanding of the molecular mechanisms that underpin intestinal homeostasis. Ageing results in a loss of intestinal epithelial homeostasis characterised by changes in architecture, cellular composition, and regenerative potential of the stem cells^1–5^. In addition, immune activation and intestinal permeability increase and there are shifts in the size and taxonomic composition of the commensal microbiota^6–15^. These phenotypes are interrelated such that interventions that target one will impact other aspects of intestinal homeostasis. Moreover, preventing a loss of intestinal homeostasis with age delays age-related decline in the organism as a whole^3,6–8,14,16–21^. Collectively, these studies have established the importance of intestinal homeostasis as a cornerstone of healthy ageing.

Alongside these studies of intestinal cell biology, physiology and microbiota we have seen growing evidence of the role of nutrition in maintaining health throughout the life course. The signalling network that regulates metabolism and growth is well accepted to influence lifelong health. This network responds to nutritional cues, and dietary nutrients and feeding patterns can themselves regulate ageing (reviewed in Longo and Anderson, 2022^22^). It is clear from this body of work that interactions between nutritional cues and longevity-associated signalling pathways are complex and context dependent. Genetics, sex, existing health status and age of onset all influence the ability of dietary interventions to extend lifespan and improve health (reviewed in Longo and Anderson, 2022^22^). Despite the challenge posed by this complexity, personalised nutrition has the potential to optimise individual health and longevity.

One key outcome of the intensive study of nutritional regulation of ageing is an appreciation that nutritional needs change significantly with age^23^ as does the ability of established nutritional interventions to extend longevity^22^. Taking this together with our knowledge of age-related intestinal decline it is surprising that few studies have addressed the impact of intestinal decline on nutrient management, a key function of the intestinal epithelium. In this regard, older animal studies suggest a reduction in transport of a range of nutrients with age^24–26^. More recently, it has been postulated that age-related defects in membrane trafficking drive loss of intestinal barrier function and might also disrupt nutrient absorption, thus underlying age-related changes in metabolic status^27^. An assumption that nutrient absorption is reduced with age pervades online sources of nutritional advice, and while these claims are not unfounded it has not yet been established that a loss of nutrient absorption is a general feature of ageing or whether such a loss impacts all nutrient types. More importantly, the mechanisms underlying such a loss, the consequences for intestinal physiology and ageing, and the impact on the effectiveness of nutritional interventions have not been explored.

A recent article summarizing the discussions that took place at an NIH workshop on diet and healthspan put forward a list of questions that will need to be considered priorities in future research, among them the question: “Do nutrient transport and uptake change with age?”^28^. However, exploring this is challenging because to date, tools enabling deeper investigation of age-related changes in intestinal nutrient management have been limited. We aim to address this in *Drosophila melanogaster*, a short-lived and genetically tractable model system that has been extensively used in studies of the ageing intestine. We present a novel assay that combines a chemically defined diet with metabolic profiling of faecal nutrient content to assess changes in nutrient uptake/excretion balance with age. Using this assay, we demonstrate that amino acid load is increased in the faecal matter of aged flies, with excess amino acids largely contributed by the microbiota. Further, age-related increase in amino acid load is associated with reduced expression of a subset of amino acid transporters in the midgut epithelium. Intestinal knockdown of the amino acid transporter *slimfast* appears protective, extending lifespan and improving microbial control in aged flies. These data highlight the contribution of age-related intestinal decline to the local nutritional environment, with consequences for health. In addition, age-related decline may drive changes in nutrient absorption capacity. Therefore, our findings have significant implications for the rational design of anti-ageing nutritional interventions.

## Results

### Drosophila Undigested Metabolite Profiling (D.U.M.P)

To measure changes in nutrient absorption/excretion balance we adopted a strategy of comparing nutritional input and output by combining a chemically defined *Drosophila* media^29,30^ with mass spectrometry profiling of faecal nutrient content. We included a non-absorbable food dye in the media to calculate the quantity of food ingested for each faecal sample by measuring its dye content. This enabled direct comparison between nutrient intake and output.

First, we needed to establish that the defined diet supported the adult stage of our wild-type Canton-S flies as expected under our laboratory conditions. Given the previously reported developmental delay of flies reared on the defined diet^29^, we reared flies on our standard cornmeal diet, transferring them to the defined diet in early adulthood. Diet is an important determinant of lifespan and age-related health and so we assessed the lifespan of our flies on the defined diet compared to our standard cornmeal diet. We found the change in diet did not alter the lifespan (Fig. S1A). We also found that flies on the defined diet continued to show an age-related increase in intestinal permeability (Fig. S1B-D) as measured by the Smurf assay^10,31^. Finally, we assessed the ability of the defined diet to support lifespan of axenic flies. We found axenic conditions resulted in lifespan extension (Fig. S1E), demonstrating that the nutritional requirements of adult female flies are fully met by the diet alone and do not need to be supplemented by the microbiota. In fact, the presence of the microbiota is detrimental with regards to lifespan in Canton-S females under our laboratory conditions.

Having confirmed the defined diet did not impact lifespan we developed a faecal collection protocol (Fig. 1A). Inclusion of FD&C blue no. 1 in the media enabled us to score “Smurf” flies that had lost intestinal barrier function and exclude them from the assay as the concentration of dye in the fecal matter is no longer representative of food intake ^10,31^. In the remaining flies the concentration of blue no. 1 in each faecal sample then provides a proxy for food consumed. Normalisation of mass spectroscopy data to dye concentration controls for differences in food intake or the quantity of faecal matter collected.

**Figure 1.**
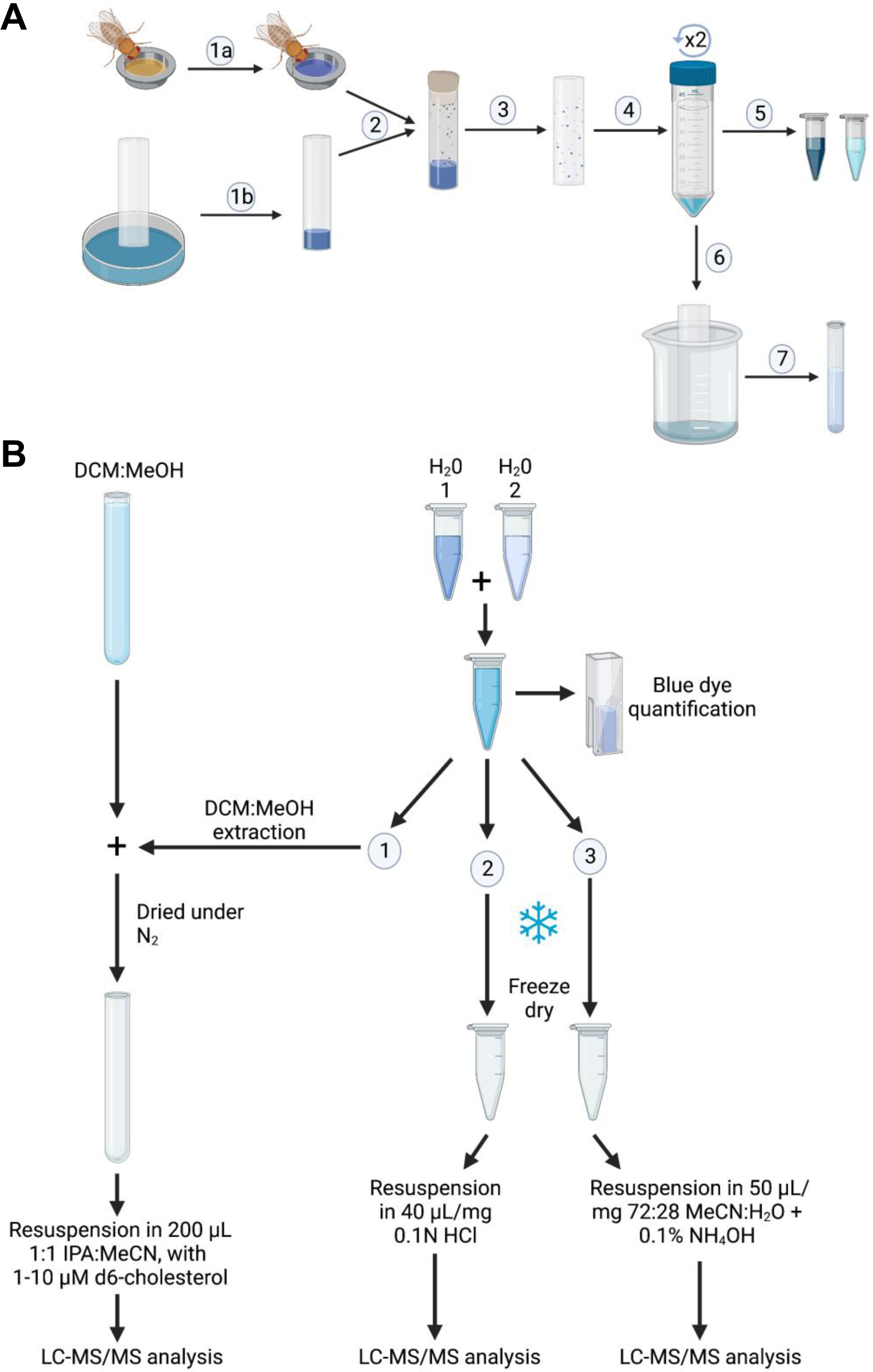
Schematic of Drosophila Undigested Metabolite Profiling assay. (A) Sample collection for the DUMP assay; 1a| Flies placed on blue food for 24h. 1b| Collection vials prepared. 2| 60 non-smurf flies transferred to collection tubes. 3| Flies incubated at 25°C for up to 5 hours, then anaesthetised and removed with food. 4| All collection vials washed within falcon tubes with the same 1.5mL H2O to produce a concentrated sample. Wash step is repeated to give a second sample 5| water samples transferred to microfuge tubes and stored at -80°C. 6| collection vials washed in 5mL 1:1 DCM:MeOH to collect non-polar fraction. 7| DCM:MeOH sample collected in glass-storage tube. Sample stored upright at - 80°C. (B) Sample processing for the DUMP assay; the two water washes are combined and 1-2 μL removed for quantification of blue dye. The remainder is split three ways for 1. Cholesterol, 2. Amino acid, and 3. Sugar analysis. DCM is added to the water fraction to extract cholesterol. The DCM layer is added to the DCM:MeOH wash sample and dried under a stream of N_2_ before resuspension 1:1 IPA:MeCN. The amino acid and sugar samples were lyophilised, then resuspended in 0.1N HCl or 72:28 MeCN:H2O + 0.1% NH4OH respectively, prior to LC-MS/MS analysis. Full details of each step are given in the materials and methods section. Created in Biorender.

For the purposes of this study, we quantified amino acids only from our final samples (see below); however, we provide details of sample preparation designed to enable measurement of amino acids, sugars and cholesterol from each faecal collection (Fig. 1B). Faecal collection involved washing faecal matter from the sides of the collection vials. Initial tests showed that two water washes were sufficient to collect 80% of amino acids and almost 50% of cholesterol (Supp. Fig. 1F). Given the time required for each wash and the diminishing returns from each additional wash, we used two water washes going forward. An additional wash with DCM:MeOH enabled collection of the remaining cholesterol (Supp. Fig. 1F), and presumably aids in collection of non-polar molecules more generally. We also found the DCM:MeOH wash to contain some amino acids, including cysteine which was not detected in the water washes. Together our faecal collection and analysis protocols result in samples that can be used to measure a wide range of nutrients and metabolites. This protocol can also be adapted for untargeted metabolomics approaches.

### Elevated amino acid levels in the faeces of aged flies originate from the microbiota

Given the importance of dietary protein levels in lifespan determination^32^ and previous reports of reduced amino acid transport with age^24^, we next applied the D.U.M.P. assay to assess changes in amino acid absorption/excretion balance with age. We found a consistent trend toward increased amino acid concentrations in faecal matter from aged flies, although not every amino acid showed a statistically significant change (Fig. 2A).

**Figure 2:**
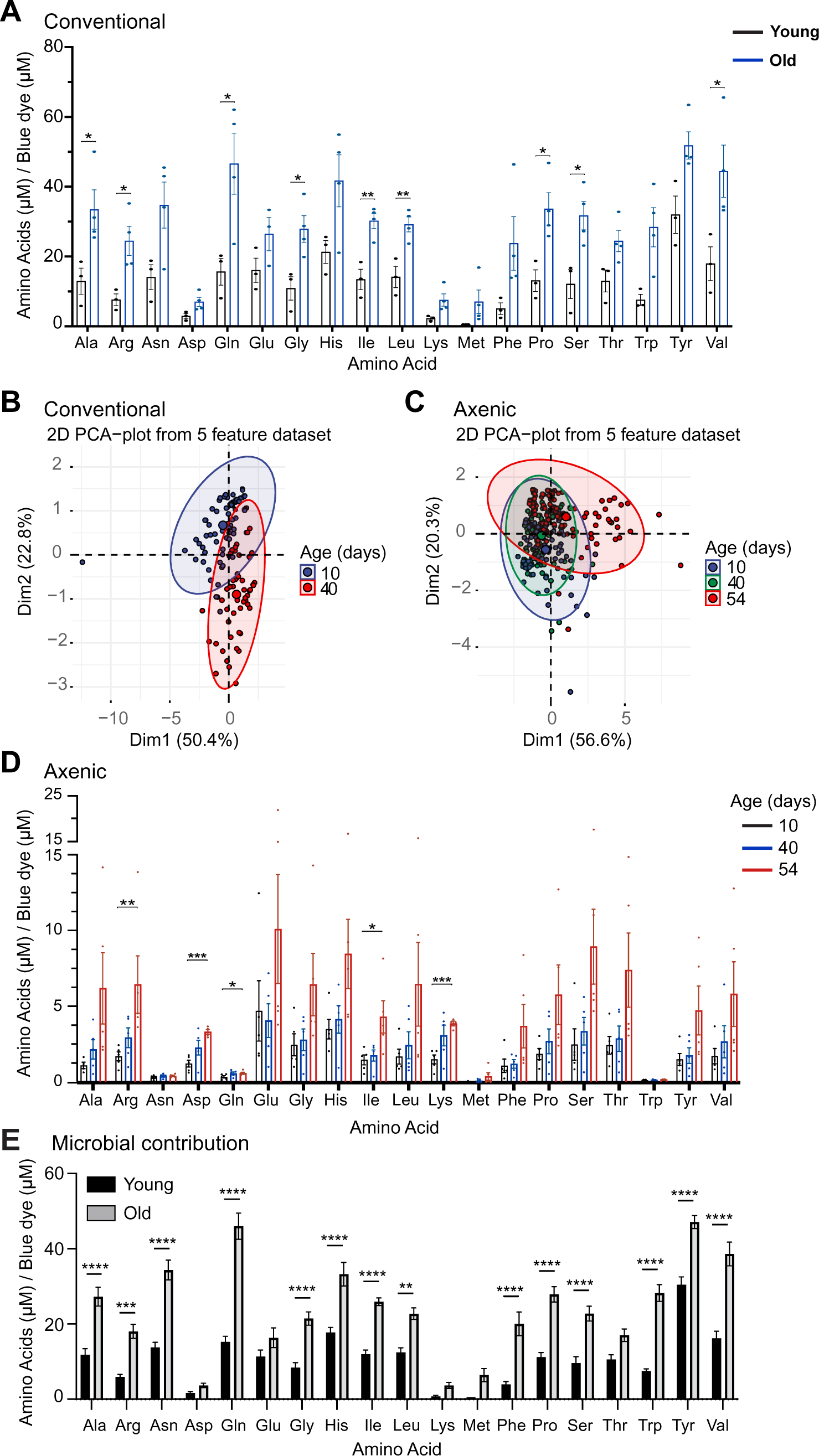
Drosophila Undigested Metabolite Profiling uncovers changes in nutrient egestion with age. (A) Relative concentration of amino acids in the faecal deposits of 10-day (9-11 day) and 40-day (39-44 day) flies, normalised to the concentration of blue food dye. (B-C) Principal component analysis for the overall structure of amino acid data from (B) conventional and (C) axenic samples plotted with respect to the first and second principal components. (D) Relative concentration in the faecal deposits of 10-day (9-11 day), 40-day (38-40 day), and 54-day (52-54 day) axenic flies, normalised to the concentration of blue food dye. Each sample pooled from >300 mated female Canton S flies. N = 3 biological replicate samples for the conventional dataset and N = 5 for the axenic data. E) Concentration of faecal amino acids, normalised to the concentration of blue food dye, contributed by the microbiota in young and old flies. The microbial contribution was calculated for each amino acid at the young timepoint by; Conventional concentration (9-11 days) – axenic concentration (9-11 days), and at the old timepoint by; Conventional concentration (38-40 days) – axenic concentration (52-54 days), using the concentration of amino acid normalised to blue dye for each sample. For each amino acid, at each time point, the calculation was carried out between every conventional and every axenic measurement. Bar graphs show mean ± SEM. Asterisks denote the results of 2-tailed unpaired T-test (normally-distributed data) or Mann-Whitney test (not-normally distributed data; (A) Trp and Try, (D) Arg, Glu, Met, Phe, Ser and Val), where * P<0.05, **P<0.01, ***P<0.001. For the axenic data tests compared 10- and 54-day samples.

We reasoned the previously reported age-related increases in microbial load in the fly intestine^6,7,10,14^ may drive this age-related increase in faecal amino acid concentrations. To test the microbial contribution to age related changes, we repeated the D.U.M.P. assay in a cohort of axenic flies. Given the lifespan extension seen in axenic flies we included a later life faecal collection. To characterise the structure of the data we performed a principal component analysis (PCA) on the conventional and axenic datasets. Plotting the data in respect to the first and second principal components, which explain the most variance, showed clustering by age in the conventional dataset (Fig. 2B). In the axenic dataset there was no separation between the 10 and 40-day faecal samples, indicating a high degree of similarity between these samples. However, the sample collected at 52-days shows separation from the 10 and 40-day samples (Fig. 2C). This is consistent with the fact that the axenic lifespan shows no increase in mortality at the 40-day timepoint, while the conventional lifespan is at 75% survival at 40 days (Supp. Fig. 1E). The axenic lifespan drops rapidly from 80% to 50% survival between days 50 and 53 (Supp. Fig. 1E), suggesting the 52-day timepoint is a closer match to the 40-day conventional sample with regards to biological age. For this reason, the 52-day sample was used for further analysis.

To determine the contribution of epithelial ageing to the changing faecal amino acid profile in the absence of the microbiota we compared our axenic samples across the age range. As suggested by our PCA analysis (Fig. 2C), a trend toward increased amino acid concentrations in the faecal matter of aged flies remained in the axenic cohort, although this was statistically significant for fewer amino acids than in the conventional dataset (Fig. 2D). The amino acids that showed statistically significant changes upon ageing were different between the conventional and axenic datasets and overall amino acid levels were much lower, demonstrating the contribution of the microbiota. Nevertheless, age-related changes in the intestinal epithelium are sufficient to drive increased amino acid load in aged fly faeces.

In order to better understand the relative contributions of epithelial changes and the microbiota to the overall aged amino acid load, we calculated the amino acid concentration contributed by the microbiota in young (10-day conventional – 10-day axenic) and old (40-day conventional – 52-day axenic) flies. This calculation was performed by comparing every conventional value to every axenic value, for each amino acid. This analysis confirmed that the microbiota is an important contributor to the faecal amino acid profile in both young and old flies and that the microbiota contributes significantly more amino acids to the faecal amino acid pool in old flies (Fig. 2E). Considering the microbial contribution as a percentage of the total amino acid load in young and old flies demonstrated that, with the exception of aspartate and lysine, the microbiota contributes over 70% of the total amino acid load (Supp. Fig. 2A). Importantly, the percentage contribution changes little with age suggesting that increased amino acid levels in aged flies are largely driven by age-related changes to the microbial population and may reflect the increased microbial load typical in aged flies.

To visualise age-related changes in microbial contribution of specific amino acids we ordered the amino acids by level of microbial contribution. Several changes in graph position are seen between young (Supp. Fig. 2B) and old (Supp. Fig. 2C) flies, confirming that changes in the microbial population contribute to age-related changes in the faecal amino acid profile as well as to amino acid levels. In addition, the order of amino acids in both young and old samples are different from that in the media (Supp. Fig. 2D), confirming the overall contribution of the microbiota to the faecal amino acid pool.

### Recapitulating aged amino acid load shortens lifespan

We assume the elevation in amino acid load in the faecal matter from aged flies reflects elevated amino acid levels in the aged intestinal lumen. We next assessed whether these changes altered lifespan. We used the D.U.M.P data to design a diet that mimics the amino acid changes that occur with age in conventionally reared flies. The amino acid concentrations in this diet, called Aged Amino Acid (AA) media, were determined by adding the difference in mean concentration between the 10- and 40-day conventional faecal samples to the concentration in the original defined media. Amino acid concentrations in the Aged AA media ranged from 1.15 to 5.18 times the concentrations in the standard defined diet. The average increase in concentration was 2.15 times that of the standard diet, equivalent to dietary protein levels that have previously been shown to reduce *Drosophila* lifespan^33^.

We found that exposure to the Aged AA media from young adult life significantly and reproducibly reduced lifespan (Fig. 3A, supp. Fig. 3A). This result was also observed in a cohort of axenic flies, suggesting it is independent of the microbiota (Fig. 3B). Consistent with this observation we found no significant impact of diet on internal bacterial load (Fig. 3C). In addition, a capillary feeding assay^34^, showed no significant difference in food intake between the standard and Aged AA media (Fig. 3D). These data are consistent with the known lifespan impact of a high protein diet^32^ and suggest that increased intestinal amino acid load may contribute to age-related decline.

**Figure 3:**
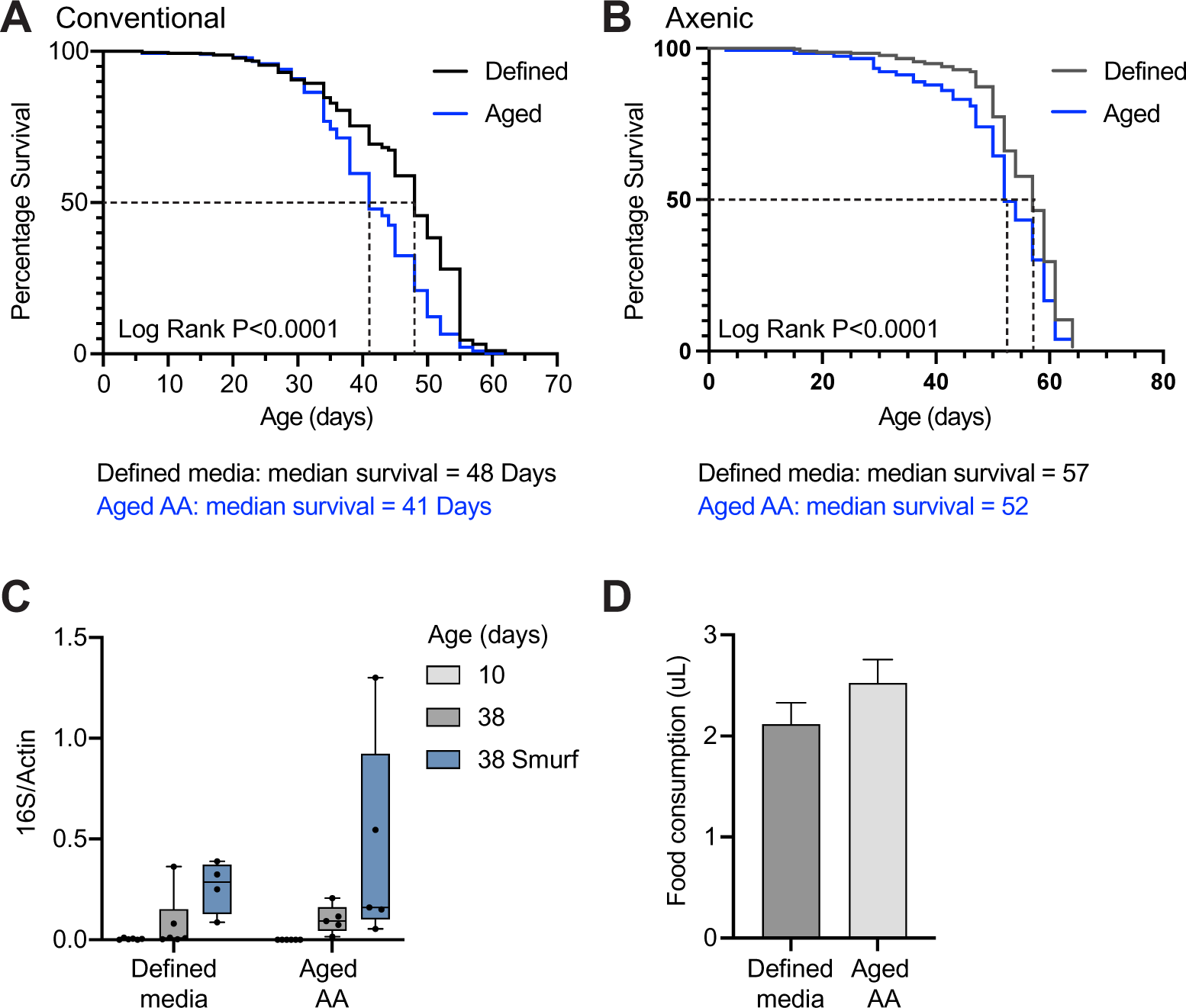
Aged AA media reduced lifespan independent of microbial status. (A) Lifespan of conventionally colonised flies on defined media or Aged AA media. Dashed black line denotes the median survival, n > 240 flies per condition at day 1. (B) Lifespan of axenic flies on defined media or Aged AA media, n > 240 flies per condition at day 1. (C) Internal bacterial load of young (10 day), old (38 day) and old smurf (38 days with intestinal barrier dysfunction) flies measured using universal primers for the 16s rRNA gene, normalised to actin. Data from at least 5 flies/sample, with a minimum of 6 samples. Box plots show the 25-75th percentiles and the median with the whiskers extending from min-max. (D) 48-hour food consumption (μL) of 9-10 groups of 10 flies fed liquid defined media or Aged AA media (omitting cholesterol). Bars show mean and standard error. No significant difference by unpaired t-test. All experiments were performed on mated Canton-S female flies.

### Expression of intestinal amino acid transporters changes with age

We next sought to understand why epithelial ageing results in increased faecal amino acid levels in the absence of the microbiota. We reasoned that this may reflect reduced amino acid transport across the intestinal epithelium with age, as previously reported for several animal species^24^. As a proxy measure of transport capacity, we assessed the expression levels of a panel of amino acid transporters in young and old intestinal tissue.

First, we generated a list of validated and predicted Drosophila amino acid transporters that contained 48 genes, 21 of which were unnamed. We selected for gut expression using online databases Flygut-*seq*^35^ and FlyAtlas 2^36^ and identified genes showing homology to human intestinal amino acid transporters using DIOPT Ortholog Finder^37,38^. This generated a final list of 18 genes spanning 8 solute carrier families (Supp. Table 1).

Of these we were able to validate quantitative PCR (qPCR) primers for 13 transporters, representing six solute carrier (SLC) families (Supp. Table 1). The validated primers were were used to quantify expression in young (10 day) and old (45 day) Canton-S female intestinal samples (Fig 4, supp. Fig 4). We found six transporter genes across 4 SLC families that showed statistically significant changes with age: sbm, Eaat1, dmGlut and slif showed significant downregulation, while CG43693 and path were upregulated (Fig 4). The amino acid substrates for CG43693 (Fig. 4A) and Sombremesa (sbm) (Fig. 4C) are unknown, and Pathetic (Path) (Fig. 4B) has been suggested to act primarily as a transceptor due to its low transport capacity^39^. However, several amino acid transporters showed decreased expression levels in aged intestines consistent with the D.U.M.P data.

**Figure 4:**
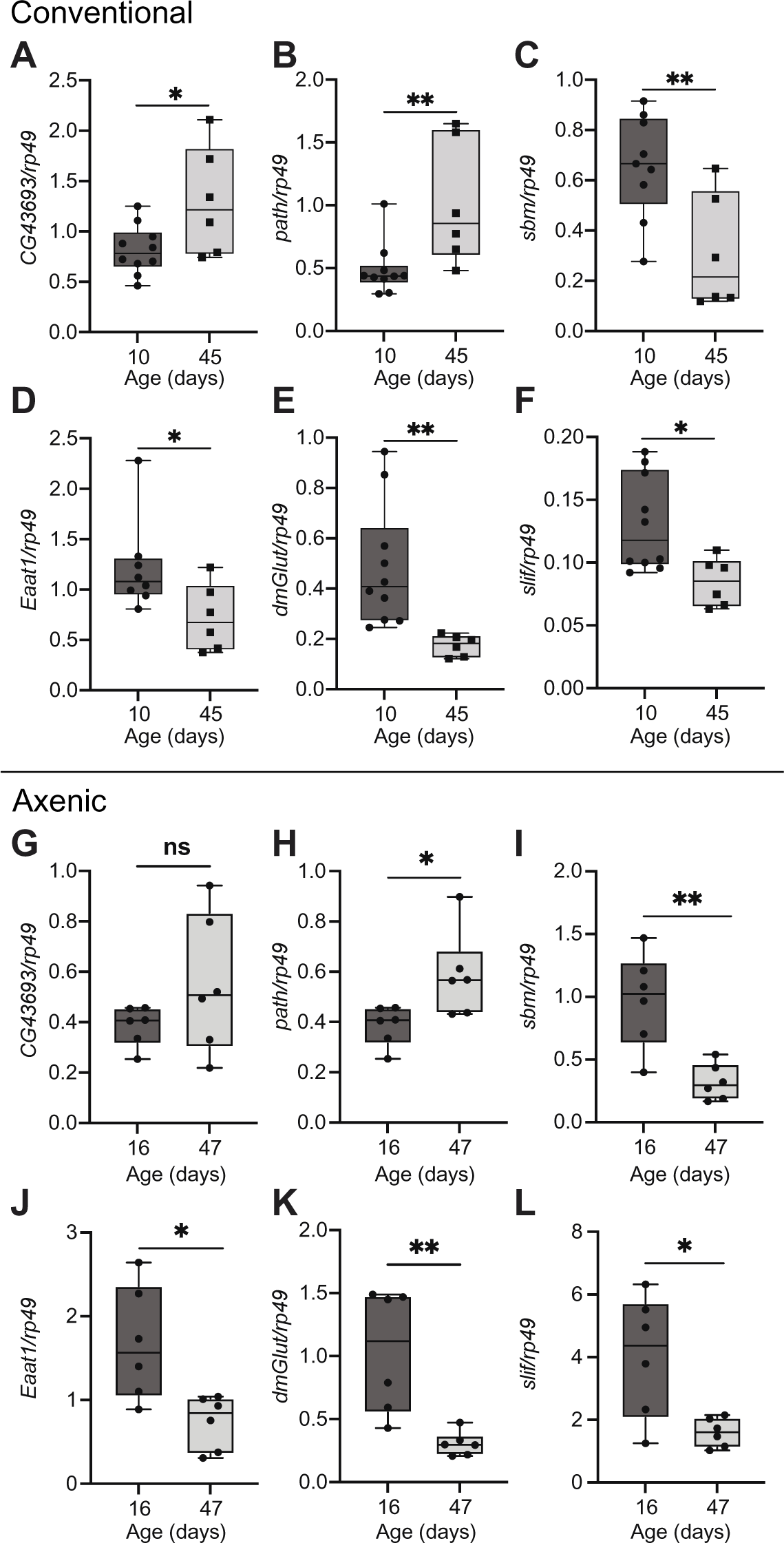
Age-related changes in amino acid transporter expression. (A-F) Transporter levels measured in dissected midguts from conventionally reared Canton-S females at 10 and 45 days of age. Normalised to the expression of rp49. N = 5 guts per sample, 8-10 samples at day 10, and 5-6 samples at day 45. (A) *CG43693*, (B) *Pathetic (path)*, (C) *sobremesa (sbm)*, (D) *Excitatory amino acid transporter 1 (Eaat1)*, (E) *dietary and metabolic glutamate transporter (dmGlut)*, (F) *Slimfast (Slif)*. (G-L) Transporter levels measured in dissected midguts from axenic Canton-S females at 16 and 47 days of age. Normalised to the expression of rp49. N = 5 guts per sample, 5-6 samples at each timepoint. (G) *CG43693*, (H) *Pathetic (path)*, (I) *sobremesa (sbm)*, (J) *Excitatory amino acid transporter 1 (Eaat1)*, (K) *dietary and metabolic glutamate transporter (dmGlut)*, (L) *Slimfast (Slif)*. Box plots show the 25-75th percentiles and the median with the whiskers extending from min-max. Asterisks denote the results of 2-tailed unpaired T-test (normally-distributed data) or Mann-Whitney test (not-normally distributed data as calculated by the Shapiro-Wilk test; *path* and *Eaat1* from conventionally reared flies (B,D)), where ns – not significant, * P<0.05, **P<0.01.

Excitatory amino acid transporter 1 (Eaat1) (Fig. 4D) transports L-glutamate with highest affinity, although L- and D-aspartate are also substrates, and there is evidence to indicate a functional role in the intestine^40^. A second glutamate transporter, dietary and metabolic glutamate transporter (dmGlut), also showed reduced expression with age (Fig. 4E). Both glutamate and aspartate show a trend toward increased levels in faecal samples from old conventionally reared and axenic flies. This increase is statistically significant for aspartate in axenic samples, suggesting the underlying cause is independent of the microbiota.

Slimfast (slif) (Fig. 4F) transports L-arginine^41^ and could be inferred to also transport lysine due to its homology to the cationic amino acid transporter family, although this has not been tested to our knowledge. A statistically significant increase in arginine and lysine levels is seen in faecal samples from old axenic flies and for arginine in faecal samples from old conventionally reared flies.

We further confirmed that the changes in expression reported for *path, sbm, Eaat1, dmGlut* and *slif* were retained in axenic flies (Fig. 4H-K). CG43693 showed a trend toward increased expression in old axenic flies, but this did not reach statistical significance. Overall, this suggests the recorded changes in transporter expression are independent of the microbiota. Taken together these data support a model whereby age-related changes in the expression levels of amino acid transporters result in reduced transport across aged intestinal epithelia which may contribute to increased luminal amino acid levels.

### Intestinal slimfast expression level regulates lifespan

We next sought to understand the physiological impact of reduced amino acid transporter expression during ageing. We focused on slimfast due to the availability of genetic tools and the significantly increased faecal concentration of its substrates with age.

To determine whether the age-related loss of *slif* expression is functionally relevant, we knocked down *slimfast* expression specifically in intestinal tissue from early adulthood. We used the *slif-antisense* line (*slif^anti^*)^41^ and the TIGS driver, a drug inducible geneswitch Gal4^42,43^, enabling us to restrict knockdown to the adult stage. We observed a significant and reproducible lifespan extension in drug induced females compared to vehicle controls (Fig. 5A and Supp. Fig 5A). Given that slimfast has been previously shown to regulate feeding^44,45^ and that reduced food intake itself can regulate lifespan we performed a capillary feeding assay^34^. We found no observable difference in the volume of food consumed (Fig. 5B). Considering that reduced intestinal *slif* may alter luminal nutrient availability, subsequently altering the intestinal microbiota, we measured bacterial load using qPCR of the bacterial 16S gene. Interestingly, we observed reduced bacterial load in old *slif^anti^* females (Fig. 5C), both prior to and following loss of intestinal barrier function. In contrast, overexpression of *slif* in the intestine for the full adult lifespan or during later adult life had no consistent impact on lifespan and did not change intestinal bacterial load (Supp. Fig. 5B-F). These data suggest loss of intestinal *slif* expression in later life may be protective, perhaps by preventing uptake of excess amino acids contributed by the aged microbiota. In addition, reduced *slif* expression may confer improved control of the intestinal microbiota, perhaps reflecting improved overall health of the aged intestine in these flies.

**Figure 5:**
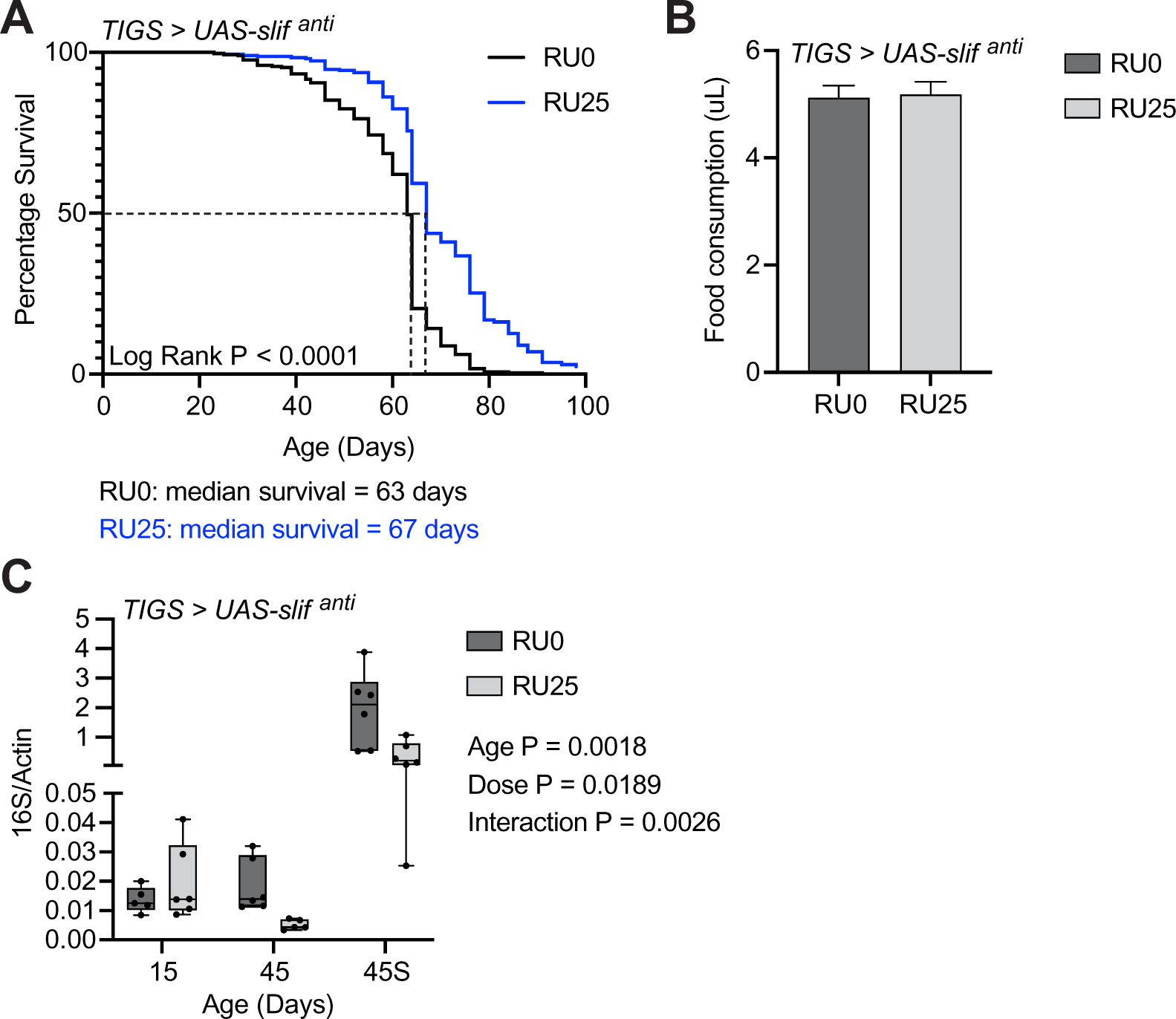
Intestinal *slimfast* levels regulate lifespan. (A) Lifespan of *TIGS > UAS-Slif^anti^* females on defined media with ethanol (RU0; black), or 25uM RU-486 (RU25; blue) throughout adult life. Dashed black line denotes the median survival. (B) 48-hour food consumption (uL) of 6 groups of 10 *TIGS > UAS-Slif^anti^* females fed liquid defined media (omitting cholesterol) for 48 hours, following 72h induction of knockdown (RU25) or control (RU0) treatment. (C) Internal bacterial load of 15 and 45 day old, and 45-day old Smurf (45S) *TIGS > UAS-Slif^anti^* females, with and without induction of intestinal *Slif* knockdown. Conventionally colonised mated females are used throughout. Bar graphs show mean and standard error. No significant difference by unpaired t-test. Box plots show the 25-75th percentiles and the median with the whiskers extending from min-max. Two-way Anova with Tukey’s multiple comparisons P-values are shown below key in panel C.

## Discussion

In this study we have developed a powerful new assay that enables us to investigate the relationship between intestinal homeostasis and nutrient management in a genetically tractable model system. We have used the D.U.M.P assay to demonstrate an increase in amino acid levels in the faecal matter, and by proxy the intestinal lumen, of aged flies. It is important to note that our targeted mass spectrometry approach measures only free amino acids. Free amino acids in the faeces will include those remaining unabsorbed from the diet, as well as those contributed by secretion from microbial or fly cells, and those released from microbial or fly proteins by digestion. Comparison between the faecal content of conventional and axenically reared flies has demonstrated the significant contribution of the intestinal microbiota to the luminal amino acid pool. This finding is in line with previous studies demonstrating the ability of the microbiota to rescue flies from undernutrition, specifically low protein diets^46,47^. In fact, we find a higher contribution to amino acid load from the intestinal microbiota than from the diet. These data also demonstrate the significant changes in this contribution that result from age-related shifts in the microbial population. The impact of changes in relative abundance of specific amino acids on health and longevity warrents further investigation, but falls outside of the scope of this study.

In addition to the direct microbial contribution, secreted proteins and shed fly cells may contribute to the nutrient pool. Elevated immune responses in the aged intestine could raise both the levels of secreted proteins (including antimicrobial peptides and other immune responsive proteins) and intestinal cell death and shedding^6,7,48,49^. These processes are also driven by age-related changes to the intestinal microbiota. Therefore, the amino acid increase recorded in the presence of the microbiota may not soley originate from microbial cells. Reduced expression levels of a subset of amino acid transporters in the aged intestine suggests that reduced amino acid transport into epithelial cells also contributes to elevated luminal levels in aged animals. Our current data cannot distinguish between amino acids remaining unabsorbed from the diet and those contributed from secreted fly proteins, shed intestinal cells or microbial cells. It will be important to address the relative contributions of these amino acid sources in future studies. However, that the percentage contribution from the microbiota remains unchanged between young and old flies suggests that changes occurring independently of the microbiota, such as reduced amino acid transport, have minimal impact on the overall age-related increase in amino acid levels.

As we would expect from previous studies^32^, increasing dietary amino acid levels shortens lifespan. In contrast, intestinal knockdown of the amino acid transporter *slimfast* extends lifespan. At first glance these data may seem contradictory, given that knockdown of *slimfast* can be expected to raise luminal levels of the amino acids that it transports. However, it is important to note that when feeding additional amino acids to young flies we assume an increased uptake of amino acids by the intestine, and therefore elevated amino acid levels in intestinal cells and other tissues. While reduced transporter expression may result in some elevation of luminal amino acid levels it will also prevent cellular uptake of excess amino acids and the resulting metabolic consequences. In effect reduced amino acid transporter expression may decouple amino acid level in the intestinal lumen from that in intestinal cells and reaching systemic circulation. Recent studies suggest elevated activity of the amino acid sensing Target of Rapamycin (TOR) pathway, an established regulator of longevity, may also have a role in intestinal decline. Recently, it has been demonstrated that TOR inhibition delays age-related intestinal decline^4,50–53^ with genetic interventions targeted to intestinal stem cells sufficient to preserve homeostasis^4,53^. Evidence is also emerging that elevated TOR activation in intestinal cells may be a feature of natural ageing^4,54^. The impact of reduced *slimfast* levels on larval growth is mediated by reduced TOR activity^41^, and we suggest that reduced *slimfast* levels in the gut may protect against elevated luminal amino acid levels via reduced TOR activity.

In this context it is also important to consider increased levels of individual amino acids, such as the *slimfast* substrate arginine, may have a very different result to an increase contributed by multiple amino acids. In fact, arginine has been shown to improve intestinal immunity phenotypes in mice, through modulation of the microbiota^55^. Increasing arginine levels also increases the diversity of murine microbiota^56,57^ improving the phenotypes of a colitis model, with benefits that are passed on through faecal transplantation and are prevented by antibiotic treatment^56,58^. In addition to impacts on the microbiota, some amino acids may have specific roles in intestinal health maintenance. For example, increased glutamate concentration has been shown to stimulate intestinal stem cell (ISC) proliferation in *Drosophila*^40^ and in mice^59^ and L-arginine has also shown to stimulate ISC expansion in mice^60^. Beyond the gut arginine has been shown to have a positive association with lifespan in a study of fibroblasts taken from a range of primate, avian and rodent species^61^ and its supplementation increases the lifespan of *C. elegans*^62^. However, arginine supplementation has also been shown to increase the secretion of IGF-1^63^ and growth hormone^64^, both of which have pro-ageing functions in mice^65–67^, and to stimulate TOR activity^68^. Increased arginine levels have been found in aged Drosophila^69^ and supplementation of 20mM arginine to the diet of Drosophila leads to a lifespan reduction, and reduced fecundity^70^. These apparent contradictions highlight the complex pleiotropic roles for individual amino acids acting in metabolism and as signalling molecules systemically and within individual tissues.

Taken together our data suggest a model whereby age-related changes to the microbiota elevate amino acid levels in the intestinal lumen while a concomitant reduction in transporter expression protects the intestinal epithelium from the metabolic consequences of excess amino acid uptake. This model has significant implications for the rational design of anti-ageing nutritional therapies. For example, while past studies have focused on the importance of dietary protein levels to healthy ageing (reviewed in Babygirija and Lamming, 2021^32^), our data highlight additional sources of elevated amino acid exposure in the aged intestine. Could reductions in dietary protein levels be effective in improving health because they counter natural increases in amino acid exposure that result from age-related intestinal decline? In addtion, there are currently calls to increase the daily recommended protein intake for elderly persons to prevent or reduce age-related muscle loss and sarcopenia^71^. Aged muscle cells are believed to be resistant to stimuli such that elderly individuals must consume higher amounts of protein to stimulate muscle growth^72–74^. However, our data suggest reduced amino acid uptake by the intestine may also underlie reduced muscle mass in the elderly. Perhaps higher quantities of protein must be consumed because a lower proportion is absorbed and bioavailable to the muscles. If this is the case, an approach that focuses on maintaining intestinal health may delay the development of sarcopenia and prevent additional risks to health resulting from high protein consumption.

In conclusion, our findings demonstrate that age-related intestinal decline profoundly changes the local nutritional environment with consequences for age-related health. While we have focused on one class of nutrient and on the impact of intestinal ageing, the D.U.M.P. assay has much broader applicability. In combination with the genetic tools available in *Drosophila* and the tractability of the *Drosophila* microbiota, the D.U.M.P. assay will enable researchers in this field to assess the impact of a wide range of perturbations on the gut nutrient pool. For example, how does nutrient availability change during infection or following a loss of cellular homeostasis and subsequent regeneration? What role does nutritional availability play in these processes? In what ways do specific microbes contribute to the nutrient pool, and how does this change following perturbation of the microbial community? Given that the local nutritional environment is not simply determined by the diet, such studies are an essential complement to efforts to understand the impact of the levels of specific nutrients on intestinal homeostasis, metabolic status, and organismal health.

## Materials and Methods

### Fly stocks

This work used the wild-type Canton-S strain (gift of Linda Partridge), TIGS-2-GeneSwitch driver line^75^ (gift of David Walker), UAS-slif^anti^ (BDSC #52655) and UAS-slif (BDSC #52661).

### Fly culture

Flies were maintained in either vials or bottles in incubators with 65% humidity and a 12-hour light/dark cycle, at either 18°C or 25°C. Stocks were maintained on cornmeal medium comprised of 1 % agar (SLS, FLY1020), 3% dried inactivated yeast (Genesee Scientific, 62-106), 1.9 % sucrose (Duchefa Biochemie, S0809), 3.8 % dextrose (Melford, G32040) and 9.1 % cornmeal (SLS, FLY1110), all given in wt/vol, and water at 90% of the final volume. Preservatives; 10 % Nipagin (SLS, FLY1136) in ethanol and an acid mixture comprised of 4.15 % phosphoric acid (VWR, 20624.262) and 41.8 % propionic acid (Sigma, P5561) in water, previously described by Lewis, 1960^76^, were added following boiling.

### Defined media

The standard defined medium was prepared as in Piper *et al*. 2014^29^ using the suggested addition of 50 % more vitamin solution and the exome matched amino acid concentrations ‘FLYAA’ reported in Piper *et al.* 2017^30^. Contrary to the published method, buffer A containing the metal ion solutions, low solubility amino acids, cholesterol, MilliQ water and acetate buffer was brought to a boil in the microwave, not autoclaved at 121 ° C for 15 minutes, except in the preparation of sterile medium for axenic flies. The remaining ingredients of essential and non-essential amino acid solutions, cysteine and glutamic acid solutions, vitamin solution, lipid-related metabolite solution, folate, 10 % nipagin in ethanol, and propionic acid were added once buffer A was cooled to 65 ° C, and thoroughly incorporated. Aged AA media was prepared in the same way, but both the quantities of amino acids used to make the stock solutions and the volume of stock solution used was altered. The masses of amino acids added to the AA media were calculated by the difference between mean old and mean young concentrations measured by the D.U.M.P. assay in conventionally reared flies, plus the original defined media amino acid concentration for each amino acid. Complete recipes for both the standard and Aged AA media are given in a supplemental protocol file attached to this manuscript.

### Lifespan

All experiments were carried out using age-matched mated females, housed at a density of 27-32 flies per vial. These flies were flipped onto new food within 24 hours of eclosing. They were then allowed at least a further 24 hours to mate before the mated females were sorted under light CO_2_ anaesthesia into vials containing defined media at 48-96 hours after eclosion. Flies were flipped to fresh food every 2-3 days and the number of dead flies was counted. At least 6 vials of 30 females were used per lifespan. To control for bacterial growth on the food, control and experimental flies are always transferred to fresh food at the same time points.

Mifepristone or RU-486 (Cayman chemicals, 10006317), hereafter referred to as RU, food was prepared by the addition of 125 µL of 20 mg/ml RU stock solution per 100mL food. To ensure sufficient volume for effective mixing 125 µL of RU stock was added to 125 µL of 100% ethanol per 100 mL of food prior to addition to the food. Control food was prepared by the addition of 250 µL ethanol per 100 mL food. Food was mixed thoroughly to ensure a homogenous solution.

### Smurf assay

The smurf assay^31^ for intestinal barrier integrity was performed by flipping flies onto food containing 2.5 % FD&C blue food dye #1 for 24 hours (FastColours, 08BLU00201). After 24 hours, the flies were flipped back onto dye-free food and the number of ‘smurf’ and ‘non-smurf’ flies were recorded.

### Generation of axenic and reassociated flies

To generate sterile (axenic) flies, eggs were collected on apple juice agar plates seeded with a paste of live bakers yeast and ddH_2_O, then washed and treated with bleach and ethanol as described previously^77^. All steps were carried out in a laminar flow hood. Briefly, <14 hour-old embryos were dechorionated in 2 % sodium hypochlorite solution for 2.5 minutes, rinsed in 70 % ethanol for 5 minutes and then rinsed with ∼200 mL sterile PBS. Embryos were then resuspended in sterile 0.01 % PBSTx (PBS + 0.01 % Triton X-100 (Sigma, X100)) and transferred to sterile autoclaved cornmeal medium.

To re-associate sterilised embryos with parental gut bacteria, 100 µL adult fly homogenate (1 fly/20 µL sterile PBS) was added to sterile cornmeal medium prior to addition of axenic embryos and allowed to dry. Flies were surface sterilised for 5 minutes in 70 % ethanol prior to homogenisation to ensure only internal microbes were present in the homogenate. Where conventionally reared flies were used as a further control to axenic or re-associated flies, 30 mated females were placed in vials containing sterilised cornmeal food and allowed to lay eggs for 24 hours, starting the evening prior to embryo collection. All medium for sterile and control flies was sterilised by autoclaving for 15 minutes at 121 °C, and all equipment used to push flies was first sterilised in Virkon for at least 20 minutes, then rinsed and stored in 80 % ethanol. Gas pads were subsequently dried by running CO_2_ through them. To maintain sterility, axenic females were separated into lifespan vials next to an open flame and flipped to new sterile food every 2-3 days in a laminar flow hood.

Sterile conditions were confirmed throughout lifespans by testing the sterility of soiled vials. Soiled stoppers were pressed into single wells of 12-well plates (Starlab, E2996-1610) containing 3 mL of pre-poured and set LB Miller Agar. The plates were incubated at 37 °C for 24-48 hours to check for bacterial growth.

### Drosophila Undigested Metabolite Profiling (D.U.M.P.) assay

#### Preparation

This assay relies on knowing the full content of the defined media. To prepare, defined media was cooked as described above, with the addition of 2.5 % blue food dye #1 after cooking (as in the Smurf assay). As well as preparing 4 mL vials, the food was poured into petri-dishes for the assembly of collection vials. Collection vials are open-ended glass tubes with 2.4 cm external diameter and 9.6 cm high, similar dimensions to standard fly vials. These were provided by the glassblowing workshop within the Durham University chemistry department. Prior to beginning the assay, collection tubes were prepared by using each tube to stamp out a disc of food from the prepared petri-dish. The food discs were secured in place in the tube ends with parafilm. This plugs one end of the vial; the other end of the tube was plugged with a foam Droso-Plug (SLS, FLY1000).

24 hours prior to sample collection, flies were flipped onto defined media containing 2.5 % FD&C blue food dye #1. This ensures that the flies’ guts will be full of the assay media containing the dye for normalisation. Additionally, this is enough time for flies losing intestinal barrier function to become clearly visible, enabling their exclusion from the assay.

#### Sample collection

To begin the assay, flies were sorted into the pre-prepared collection vials under light CO_2_ anaesthesia. Approximately 55 - 65 non-smurf, mated females were sorted into each vial. More flies do not increase the amount of faecal deposit as when overcrowded they get stuck at the bottom of the vial. 6 collection vials were used per sample to ensure a large enough sample for full analysis. Once the flies awoke from anaesthesia, they were incubated at 25°C for between 4 and 5 hours to eat and defecate as usual. Standing the vials at a 45° angle results in an improved sample, particularly when collecting samples from aged flies with reduced mobility. Following incubation, the flies were anaesthetised with CO_2_, the parafilm removed and the food disc along with the flies was pushed out from the bottom of the tube. The outside of the vial and the food residue was cleaned with a tissue dampened by milliQ water until the tissue came away clean.

The collection vials were all the same weight (26.91g ± 0.41) and fit inside 50mL falcon tubes. Once within the 50 mL tube, the collection vials were thoroughly rinsed with 1.5 mL of water to collect the polar, water-soluble faecal fraction, and spun down. The same 1.5 mL of water was used to rinse each tube. This ensured the sample could be frozen and lyophilised quickly. Two falcon tubes are used, with the collection vials each being washed, then the water sample split between the falcons, to ensure the tubes are balanced for centrifugation. Once all tubes are rinsed, the sample is transferred to a microcentrifuge tube. The rinse is repeated a second time to ensure all faecal matter is collected, resulting in two water-wash samples of 1.5 mL.

The vials were then rinsed thoroughly with a 1:1 mixture of DCM and methanol to collect non-polar deposits. The rinses were carried out in a small glass beaker to collect the sample. The volume for this wash was not limited, but approximately 5 mL was used for all the tubes. Plastic use should be avoided with the DCM:MeOH wash. All samples were frozen at -80 ° C prior to downstream processing.

#### Sample processing

The two water washes were combined, and 1-2 μL was removed and diluted 1/100 for the blue dye to be quantified at 628nm absorbance, on the Bio-tek Synergy H4, alongside a standard curve of known concentration. The remaining sample was split three ways to enable analyte-specific processing and quantification. While we only quantified amino acids in our final samples for the purpose of this study, we provide details for initial processing of the other sample types here. The amino acid and sugar fractions were lyophilised and weighed, then resuspended in 40 μL 0.1N HCl, or 72:28 MeCN:H_2_O + 0.1% NH_4_CN per mg respectively. The resuspended amino acid sample was diluted 1:100 in HILIC A for LC-MS/MS analysis, and the sugar samples were diluted between 1:10 and 1:100 in the 72:28 MeCN:H_2_O + 0.1% NH_4_CN prior to LC-MS/MS analysis. The third fraction was for cholesterol analysis. An equal volume of DCM was added to the water fraction, then the sample was vortexed for 2 minutes prior to 15 minutes of centrifugation at 1000x rpm. The DCM layer was removed and added to the DCM:MeOH wash, this was dried under a stream of N_2_, then resuspended in 200 μL 1:1 IPA:MeCN, and spiked with 10 μM d6-cholesterol prior to LC-MS/MS analysis.

#### Amino acid quantitation

LC-MS/MS analysis was performed on a Shimadzu Nexera X2 Ultra-Fast Liquid Chromatography system consisting of binary pumps, an on-line degassing unit, an autosampler, and a column oven (Shimadzu Corporation, Kyoto, Japan), coupled with an AB Sciex 6500 QTRAP mass spectrometer consisting of an electrospray ionization (ESI) source (AB SCIEX, Framingham, MA, USA). 1uL of sample of injected on to a BEH Amide (1.0 x 100mm, Waters, Milford, MA, USA), BEH HILIC (1.0 x 100mm, Waters, Milford, MA, USA) or Ascentis HILIC (2.1 x 150mm, Waters, Milford, MA, USA) maintained at 40 °C, at a flow rate of 0.2 mL/min or 0.3mL/min for the 1.0 and 2.1mm columns respectively. Three columns were used to enable separation of isomers leucine and isoleucine, and optimal peak shape for more accurate quantification. The mobile phases consisted of Solution A (85% MeCN + 0.15% FA + 10mM ammonium formate) and Solution B (85% H_2_) + 15% MeCN + 0.15% FA + 10mM ammonium formate). A gradient elution was optimized for the separation of amino acids with 1.5 min equilibration 0-6.0 min, 0% Solution B; 6.0-10 min, 0-100% Solution B; 10-12 min, 100% Solution B; 12-12.1 min, 0% Solution B; 12.1-18 min, 0% Solution B; with a total run time of 21 min. The ion source was operated in positive mode under an optimal condition: curtain gas, 40 psi; nebulizer gas, 50 psi; auxiliary gas, 50 psi; ion spray voltage, 5500 V; and temperature, 500°C. Optimal multiple-reaction monitoring (MRM) transitions were identified for the amino acids (Supplemental table 2). Quantification was performed using an external calibration curve, amino acids stocks in 0.1N HCl were diluted with HILIC A as per sample preparation. Data acquisition and analysis were all performed with Analyst 1.6.3 software (AB SCIEX)

#### Analysis

Relative quantification was performed using the blue food dye; the mass-spec calculated concentration (μM) was adjusted for the sample dilution. This was then divided by the concentration of the blue dye (μM) for that sample.

The percentage change between young and old timepoints was calculated using the equation:

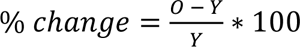

Where O is the normalised concentration at the old timepoint, and Y is the normalised concentration at the young timepoint. The young concentrations from all replicates were compared against the old concentrations from each replicate.

The microbial concentration was calculated at young and old timepoints by:

*Microbial contribution = conventional concentration − axenic concentration*

Where the conventional concentrations from each replicate were compared against the axenic concentrations for each replicate and at every timepoint.

### Capillary feeding (CAFÉ) assay

Food consumption was measured, essentially as previously described^34^, using liquid defined media made without agar and, due to issues in solubility, without cholesterol. Cotton wool balls were placed at the bottom of drosophila vials and saturated with water. Any excess water was removed, and the vial walls were dried. 10 μL pipette tips were trimmed to enable capillary tubes to extend from the end with a tight fit so they wouldn’t slip. Drosophila flugs were cut in half laterally, and three prepared tips were inserted through the flug. The capillary tubes were prepared by loading with 2 μL of mineral oil to limit evaporation, and 10 μL of liquid food, then the interface between oil and food was marked.

10 mated female flies were sorted into the prepared vials. Each condition had a minimum of 10 vials. Additionally, a minimum of three evaporation control vials, containing no flies, were used for each experiment. Once the flies were awake within the vial, the capillary tubes were inserted into the tips, so the ends extended just beyond the base of the tip. The capillary tubes were changed every 24 hours, and the distance between the interface between oil and food, and the mark denoting the starting interface was recorded. The flies were allowed 24 hours to acclimate to the new feeding setup prior to measurements being recorded. Following acclimation, the assay was performed for 24-48 hours. Food consumption was calculated in mm, by calculating the difference in the start and end interface and subtracting the average difference in the start and end interface of the evaporation controls. A conversion factor (X μL/mm) was calculated for each capillary tube by dividing 10 (the amount of liquid food), by the starting interface (mm). This was then used to convert food consumption in mm to μL.

### Bacterial DNA extraction

30 whole flies were sorted under light CO_2_ anaesthesia into microcentrifuge tubes and frozen at -80 °C. Flies were then surface sterilised in 70 % ethanol for 5 minutes and rinsed twice in sterile PBS. 5 flies were sorted into sterile microcentrifuge tubes and homogenised in 200 µl lysis buffer (1 X TE, 1 % Triton X-100, 1/100 proteinase K (NEB, P8107S)) using a mechanical pestle, with a further 300 µl of lysis buffer being added after homogenisation. Lysate was incubated for 1.5 hours at 55 °C then transferred to screw cap vials containing 0.4 g sterilised low binding 100-micron silica beads (OPS Diagnostics, BLBG 100-200-11) and vortexed for 10 minutes at max speed. Lysate was then transferred back to the original tube and incubated for a further 1.5 hours at 55 °C followed by 10 minutes at 95 °C.

### RNA extraction

Total RNA was extracted from 5 midguts per biological replicate. Midguts were dissected in ice-cold PBS and included the cardia and midgut, through to the midgut-hindgut junction, excluding the hindgut and Malpighian tubules. Midguts were homogenised in 100 µL TRIzol™ (Thermofisher, 15596026) using a motorised pestle. RNA was extracted following the standard TRIzol protocol.

### cDNA preparation

For a 10 µL reaction, 1 µL DNase buffer (Thermo, EN0521) and 0.5 µL DNase 1 (Thermo, EN0521) were added to 8.5 µL RNA solution and incubated for 30 minutes at 37 °C, to remove any DNA contaminants. The DNase enzyme was then deactivated by the addition of 0.5 µL 50mM EDTA (Thermo, EN0521) and subsequent incubation for 10 minutes at 65 °C. cDNA synthesis was carried out using Thermo Scientific’s First Strand cDNA Synthesis kit. In short, 1 µL of random hexamers (Thermo, SO142) were added and the samples were incubated at 70 °C for 5 minutes before a 9 µL of a master mix containing 4 µL first strand buffer (Thermo, EPO442), 0.5 µL RIbolock RNase inhibitor (Thermo, EO0382), 2 µL 10mM dNTPs (Thermo, R0181), 0.2 µL RevertAid reverse transcriptase (Thermo, EPO442) and 2.3 µL molecular grade water (Sigma, W4502) per well, was added. The samples were then incubated for 10 minutes at room temperature, followed by 1 hour at 37 °C and 10 minutes at 70 °C. The samples were spun following each incubation step.

### Quantitative PCR

qPCR was performed with Power SYBR™ Green master mix (Fisher, 10219284) on a Bio-RAD CFX Connect Real-Time PCR detection system. Either 4.5 µL bacterial DNA or 1 µL of *Drosophila* cDNA sample was used in a 10 µL reaction with 5 µL of Power SYBR™ Green and 0.5 µL of L + R primer mix. Any remaining volume was made up with molecular grade water. Cycling conditions were: 95°C for 10 minutes, then 40 cycles of 95°C for 15 seconds followed by 60°C for 60 seconds. All calculated gene expression values were normalised to the value of the housekeeping gene rp49. Levels of the bacterial 16S rRNA gene were normalised to the level of the *Drosophila* Actin gene. The primer sequences used are given in table 1.

**Table 1:**
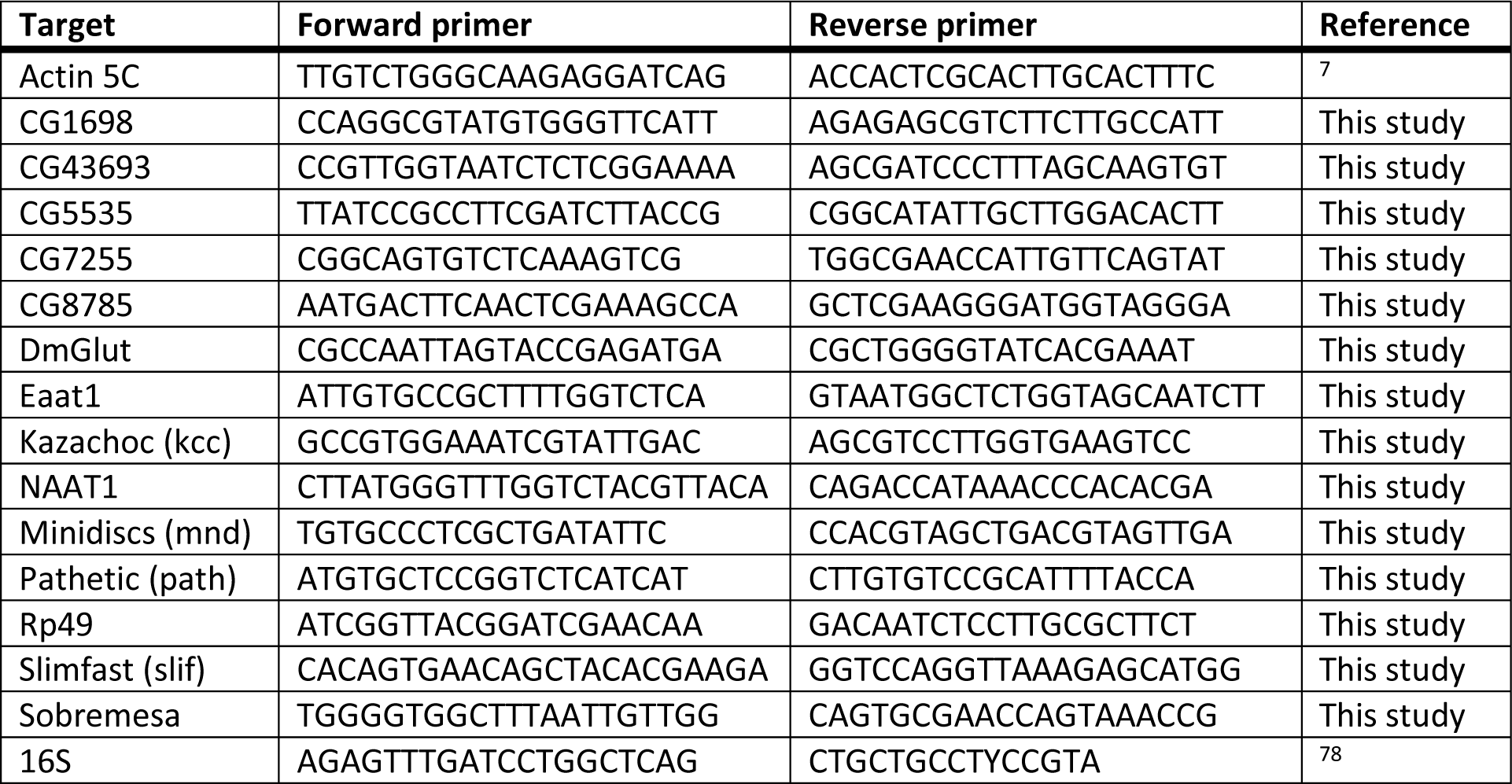
Primer sequences.

### Statistics

All Statistical analyses were performed using GraphPad Prism 9, excluding the binomial tests and principal component analyses which were performed in R (Rstudio build 372, R version 4. 0. 3). Survival curve analysis was completed using the log-rank test in GraphPad Prism; the median survival was calculated by Graph Pad Prism 8 and is an approximation to the closest sampling time. Quantitative PCR data and normalised DUMP amino acid data were cleaned in GraphPad Prism, with outliers detected using the Robust regression and outlier removal (ROUT) method with Q=1. Normality of the cleaned data was tested by Shapiro-Wilk tests to determine the most appropriate statistical test. Differences between two points were analysed by students t-test, or mann-whitney tests for normally and non-normally distributed data respectively. Differences in gene expression values between more than two time points, within or between conditions, were analysed by *two way Anova with Tukeys multiple comparison.* P-values less than 0.05 were considered statistically significant. Bar-graph error bars depict ± standard error of the mean (SEM). For all box-plots shown, boxes display the 25-75^th^ percentiles, with the horizontal bar at the median, and whiskers extending from the minimum to maximum points.

## Supporting information

Supplemental defined media protocol

## Acknowledgements

We thank L. Partridge, D. Walker, and the Bloomington Drosophila stock center for fly stocks, H. Siddle for technical support, the glass workshop at Durham University for providing collection vials, Jeanette Alcaraz, Anona Galbraith and Sophie Waldron for helpful discussions. This work was supported by the BBSRC doctoral training program and Durham University. Stocks obtained from the Bloomington Drosophila Stock Center (NIH P40OD018537) were used in this study.

---

**Supplemental figure 1.**
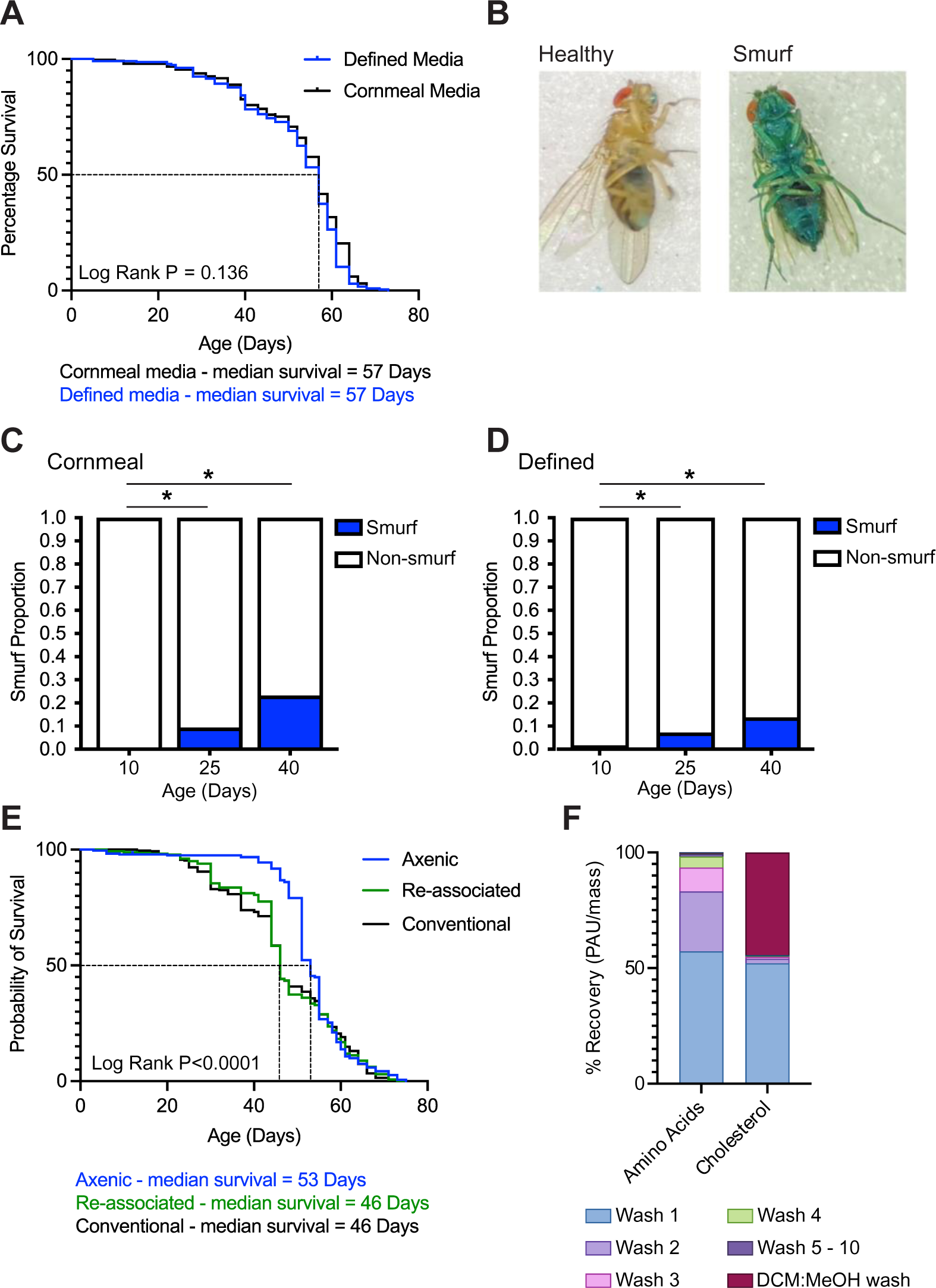
Defined media does not impact lifespan. (A) Lifespan of Canton-S female flies fed during adulthood on cornmeal or defined media. N > 200 per condition at day 1 (B) Images displaying the characteristic smurf phenotype. (C-D) Proportion of smurf flies within the total population taken at the ages shown for (C) cornmeal and (D) defined media. N > 200 per condition at day 1 (E) Lifespan of axenic, re-associated and conventionally reared flies fed during adulthood on defined media. N > 450 Canton-S females per condition at day 1. Asterisks denote the result of binomial tests, where * p<0.05. (F) Percent recovery; the sum of percent abundance (PAU)/mass of all amino acids or cholesterol per wash, as a percentage of the sum of PAU/mass of all amino acids or cholesterol for all washes. No cholesterol was detected following wash 5.

**Supplemental figure 2:**
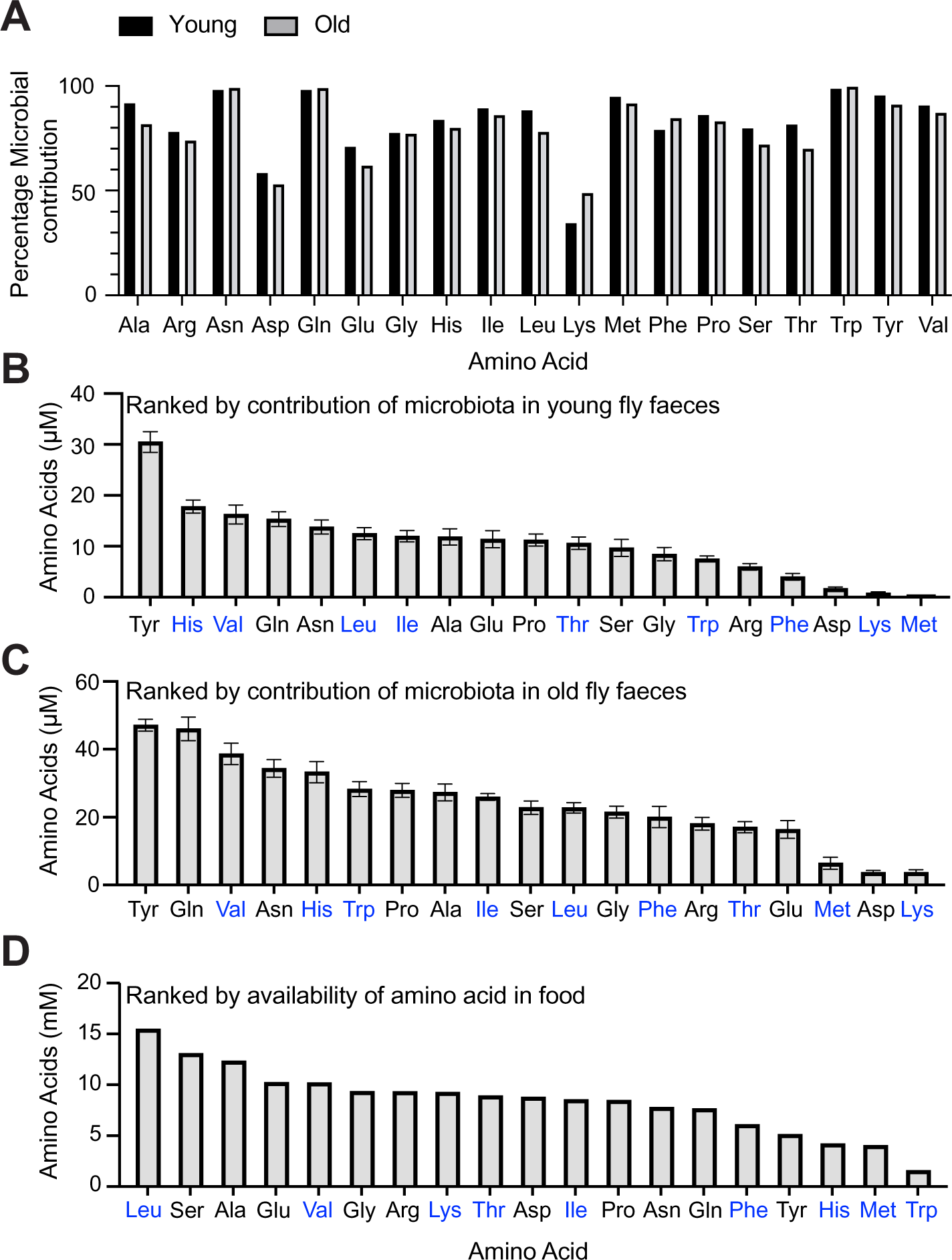
Microbial contribution with age shifts faecal amino acid profile. (A) The mean microbial contribution (Fig. 2E) shown as a percentage of the mean total amino acid load (Fig. 2A) for each amino acid in both young and old flies. (B-D) Amino acids ordered by level of microbial contribution in (B) young and (C) old flies, and by (D) availability within the food. Blue amino acid labels denote essential amino acids. Bar graph shows mean ± SEM. Asterisks denote the results of a 2-way Anova, where * P<0.05, **P<0.01, *** P<0.001, ****P<0.0001

**Supplemental Figure 3:**
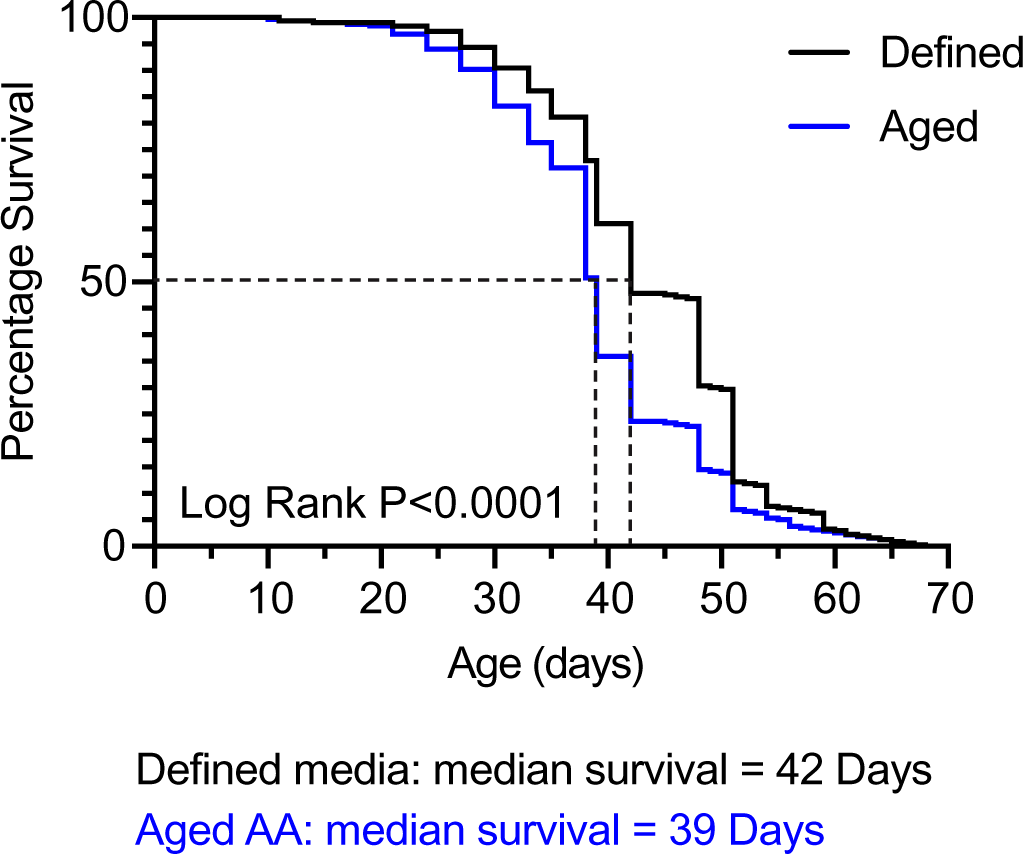
Aged AA media reduced lifespan. An independent replicate lifespan of conventionally colonised mated Canton-S females on defined media or Aged AA media. Dashed black line denotes the median survival, n > 240 flies per condition at day 1.

**Supplemental Figure 4:**
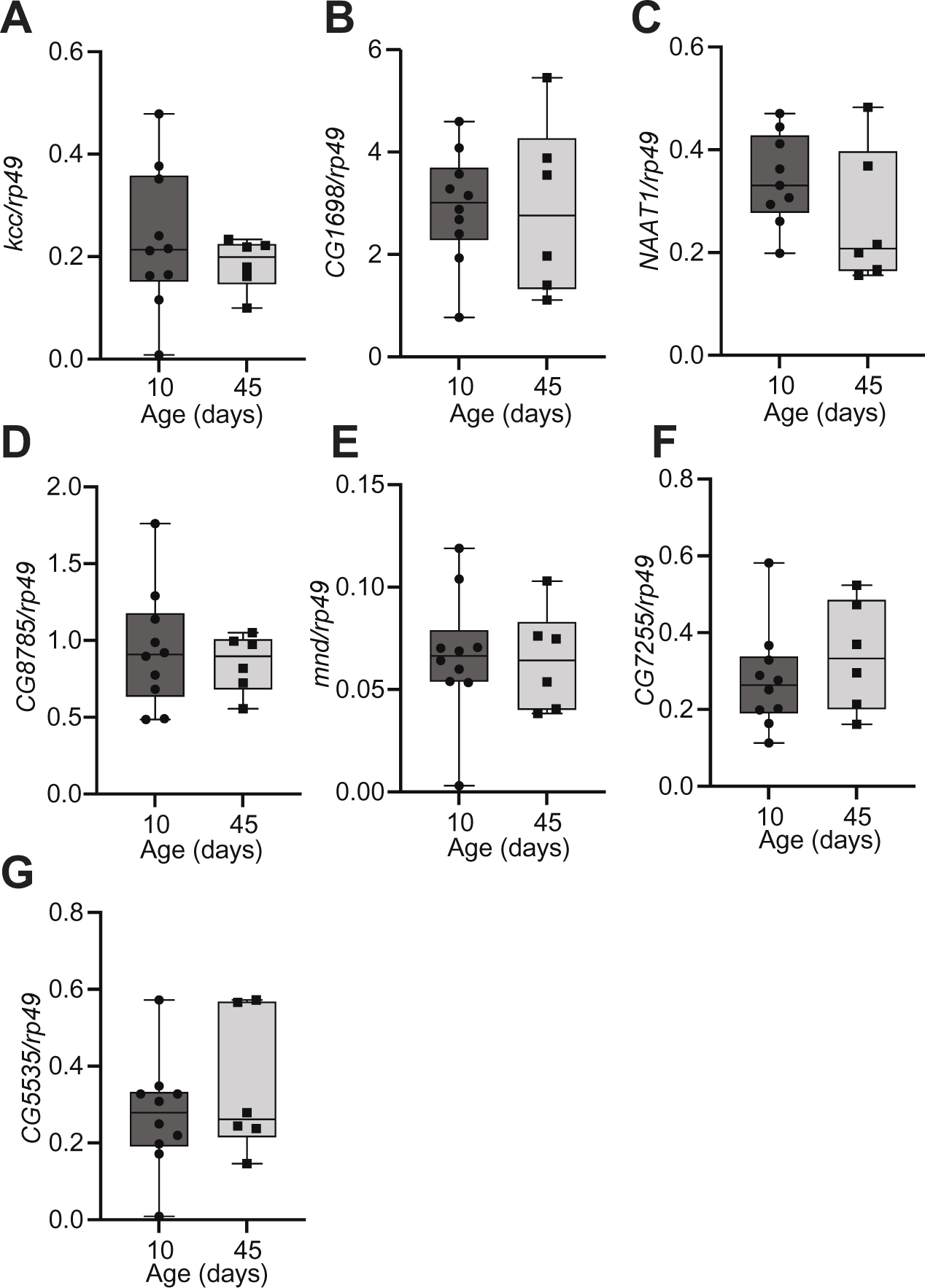
Some amino acid transporters do not show changes in expression with age. Transporter levels measured in dissected midguts from conventionally reared Canton-S females at 10 and 45 days of age. Normalised to the expression of rp49. N = 5 guts per sample, 8-10 samples at day 10, and 5-6 samples at day 45. (A) *kazachoch (kcc),* (B) *CG1698*, (C) *Nutrient amino acid transporter 1 (Naat1),* (D) *CG8785,* (E) *minidiscs (mnd),* (F) *CG7255,* and (G) *CG5535.* Box plots show the 25-75th percentiles and the median with the whiskers extending from min-max.

**Supplemental Figure 5:**
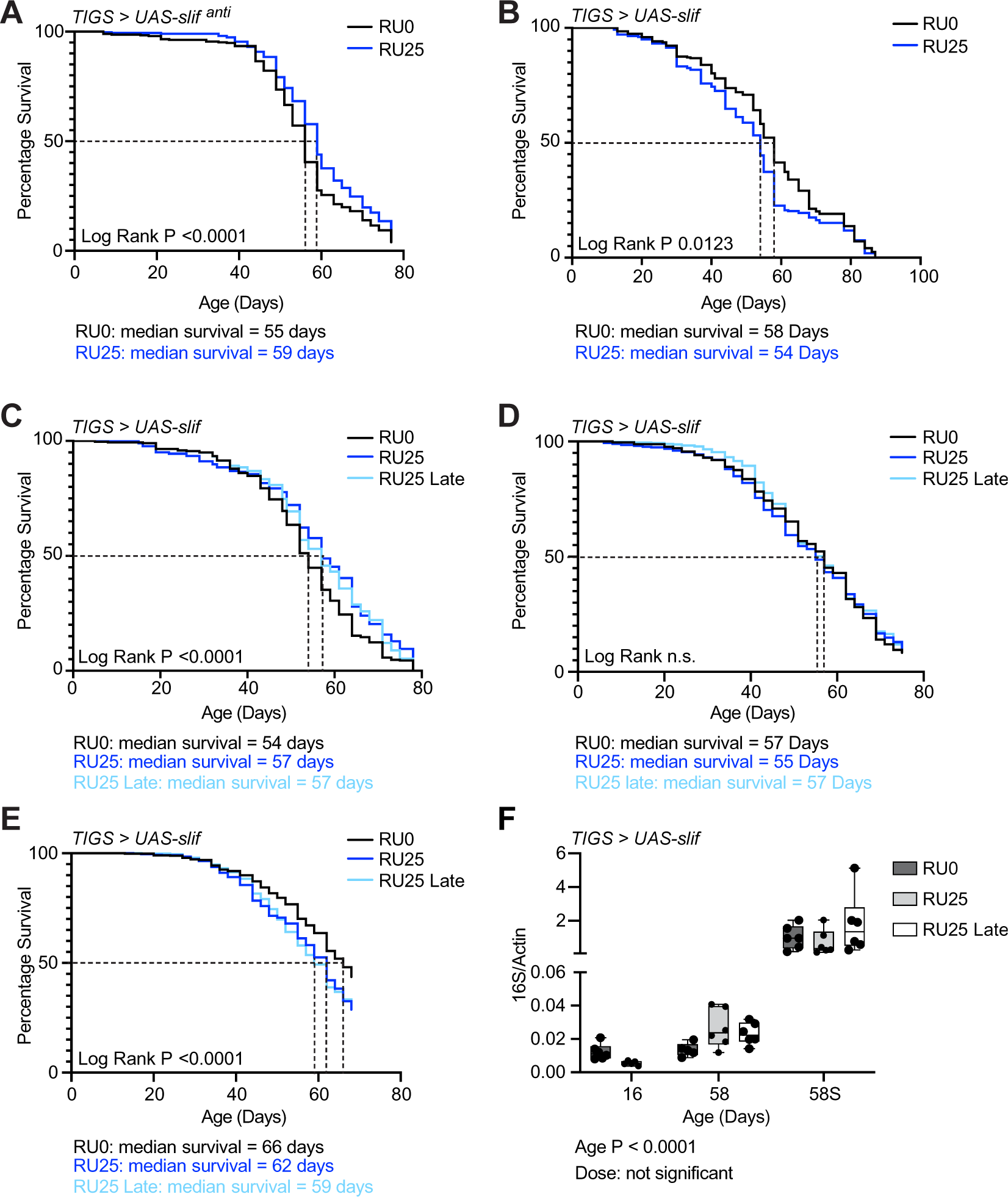
Intestinal *slimfast* levels regulate lifespan. (A) Replicate lifespan of conventionally colonised mated *TIGS > UAS-slif^anti^* females on defined media with ethanol (RU0; black), or 25uM Ru-486 (RU25; blue) throughout adult life. Dashed black line denotes the median survival. (B-E) Replicate lifespans of conventionally colonised mated *TIGS > UAS-slif* females on defined media with ethanol (RU0; black), 25uM Ru-486 throughout adult life (RU25; dark blue) or from day 35 (RU25 late, pale blue). Dashed black line denotes the median survival. (F) Internal bacterial load of 16 and 58 day old, and 58-day old Smurf flies (58S) mated female *TIGS > UAS-slif* females, with induction of intestinal *slif* overexpression throughout adult lifespan (RU25) or from day 35 (Late RU25) and ethanol treated controls (RU0). Box plots show the 25-75th percentiles and the median with the whiskers extending from min-max. Two-way Anova with Tukey’s multiple comparisons P-values are shown below graph in panel F.

**Supplemental Table 1:**
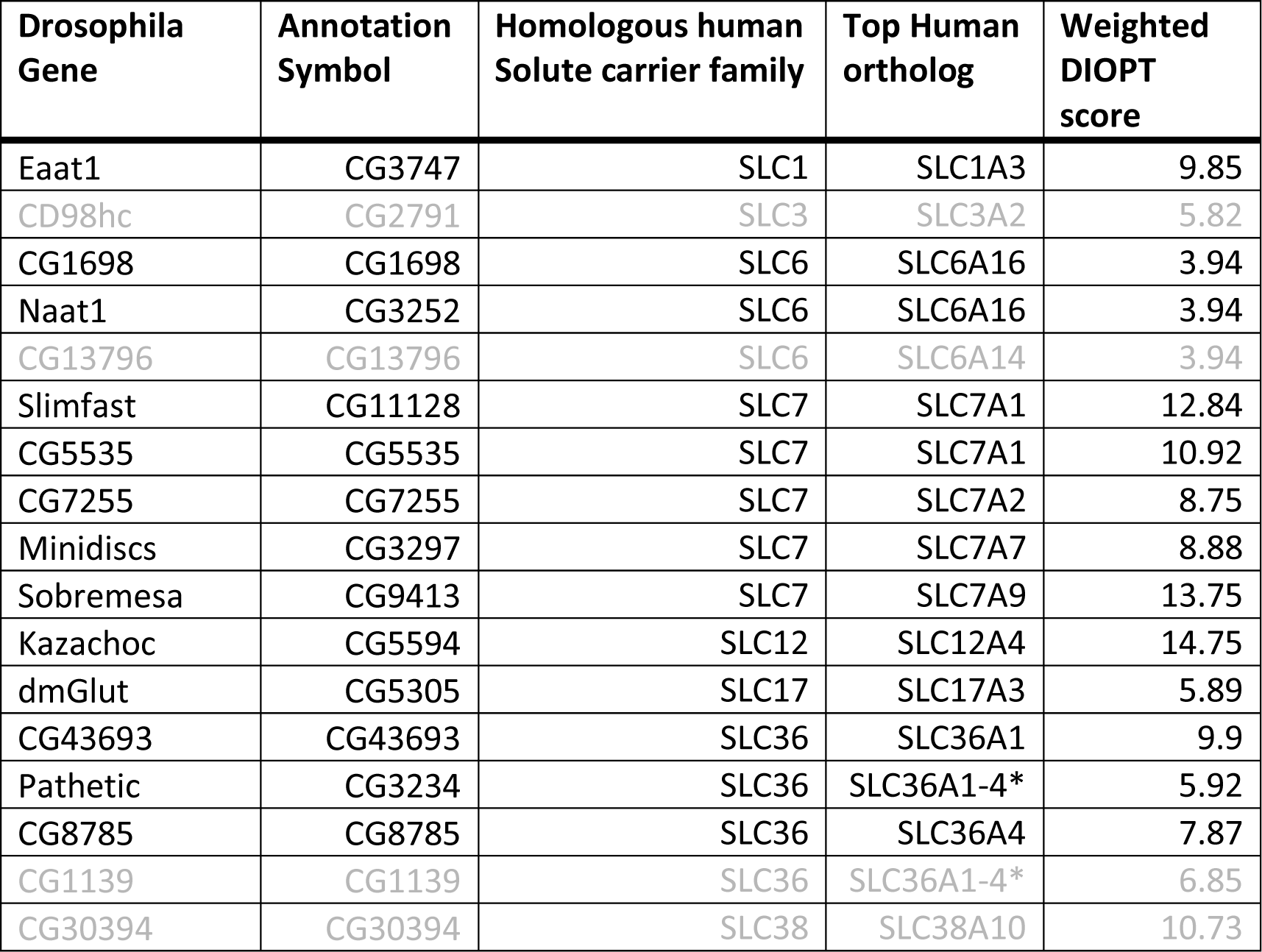
Final list of genes with validated or predicted amino acid transporter function and predicted intestinal expression. Top human orthologs and weighted DIOPT score determined from https://www.flyrnai.org/cgi-bin/DRSC_orthologs.pl. The genes in grey font were not taken forward due to poor qPCR primers. *All top orthologs, sharing the same weighted DIOPT score.

| Drosophila Gene | Annotation Symbol | Homologous human Solute carrier family | Top Human ortholog | Weighted DIOPT score |
| --- | --- | --- | --- | --- |
| Eaat1 | CG3747 | SLC1 | SLC1A3 | 9.85 |
| CD98hc | CG2791 | SLC3 | SLC3A2 | 5.82 |
| CG1698 | CG1698 | SLC6 | SLC6A16 | 3.94 |
| Naat1 | CG3252 | SLC6 | SLC6A16 | 3.94 |
| CG13796 | CG13796 | SLC6 | SLC6A14 | 3.94 |
| Slimfast | CG11128 | SLC7 | SLC7A1 | 12.84 |
| CG5535 | CG5535 | SLC7 | SLC7A1 | 10.92 |
| CG7255 | CG7255 | SLC7 | SLC7A2 | 8.75 |
| Minidiscs | CG3297 | SLC7 | SLC7A7 | 8.88 |
| Sobremesa | CG9413 | SLC7 | SLC7A9 | 13.75 |
| Kazachoc | CG5594 | SLC12 | SLC12A4 | 14.75 |
| dmGlut | CG5305 | SLC17 | SLC17A3 | 5.89 |
| CG43693 | CG43693 | SLC36 | SLC36A1 | 9.9 |
| Pathetic | CG3234 | SLC36 | SLC36A1-4* | 5.92 |
| CG8785 | CG8785 | SLC36 | SLC36A4 | 7.87 |
| CG1139 | CG1139 | SLC36 | SLC36A1-4* | 6.85 |
| CG30394 | CG30394 | SLC38 | SLC38A10 | 10.73 |

**Supplemental table 2.** MRM transitions for the amino acids quantified.

| Q1 m/z | Q3 m/z | Time (msec) | Amino Acid | DP (V) | EP (V) | CE (V) | CXP (V) |
| --- | --- | --- | --- | --- | --- | --- | --- |
| 166.1 | 120.1 | 50 | Phe 1 | 30 | 10 | 15 | 12 |
| 132.1 | 86.1 | 50 | Leu 1 | 36 | 10 | 23 | 14 |
| 132.1 | 86.1 | 50 | Ile 1 | 40 | 7 | 15 | 8 |
| 150 | 104 | 50 | Met 1 | 30 | 10 | 15 | 12 |
| 118.1 | 72.1 | 50 | Val 1 | 30 | 10 | 15 | 12 |
| 116 | 70.1 | 50 | Pro 1 | 30 | 10 | 15 | 12 |
| 182.1 | 136.1 | 50 | Tyr 1 | 30 | 10 | 15 | 12 |
| 90.1 | 44.1 | 50 | Ala 1 | 30 | 10 | 10 | 12 |
| 120 | 74.1 | 50 | Thr 1 | 30 | 10 | 15 | 12 |
| 76 | 30.1 | 50 | Gly 1 | 46 | 10 | 20 | 12 |
| 106 | 60 | 50 | Ser 1 | 30 | 10 | 10 | 12 |
| 148 | 84 | 50 | Glu 1 | 30 | 10 | 15 | 12 |
| 134 | 88 | 50 | Asp 1 | 26 | 10 | 13 | 14 |
| 156 | 110.1 | 50 | His 1 | 30 | 10 | 15 | 12 |
| 175.1 | 70.1 | 50 | Arg 1 | 30 | 10 | 15 | 12 |
| 147.1 | 84.1 | 50 | Lys 1 | 30 | 10 | 15 | 12 |
| 166 | 103 | 50 | Phe 2 | 31 | 10 | 35 | 28 |
| 132.1 | 69 | 50 | Leu 2 | 36 | 10 | 23 | 20 |
| 150 | 133 | 50 | Met 2 | 21 | 10 | 13 | 8 |
| 118 | 55 | 50 | Val 2 | 36 | 10 | 29 | 6 |
| 182.1 | 165 | 50 | Tyr 2 | 46 | 10 | 19 | 6 |
| 120 | 56 | 50 | Thr 2 | 86 | 10 | 23 | 6 |
| 106 | 42 | 50 | Ser 2 | 31 | 10 | 31 | 4 |
| 148 | 102 | 50 | Glu 2 | 21 | 10 | 15 | 4 |
| 134 | 70 | 50 | Asp 2 | 36 | 10 | 25 | 6 |
| 156 | 93 | 50 | His 2 | 41 | 10 | 31 | 8 |
| 175 | 116 | 50 | Arg 2 | 46 | 10 | 19 | 6 |
| 147 | 130 | 50 | Lys 2 | 16 | 10 | 13 | 6 |
| 122 | 76 | 50 | Cys 1 | 30 | 10 | 10 | 15 |
| 205.1 | 146.1 | 50 | Trp 1 | 36 | 10 | 10 | 23 |
| 147.1 | 84.1 | 50 | Gln 1 | 30 | 10 | 10 | 15 |
| 133 | 74 | 50 | Asn 1 | 30 | 10 | 25 | 25 |
| 133 | 116 | 50 | Asn 2 | 31 | 10 | 13 | 8 |
| 205.1 | 188 | 50 | Trp 2 | 41 | 7 | 15 | 9 |
| 147 | 102 | 50 | Gln 2 | 31 | 10 | 19 | 12 |
| 132.1 | 69 | 50 | Ile 2 | 40 | 7 | 24 | 9 |

## References

1. Choi, N.-H., Kim, J.-G., Yang, D.-J., Kim, Y.-S. & Yoo, M.-A. Age-related changes in Drosophila midgut are associated with PVF2, a PDGF/VEGF-like growth factor. Aging Cell 7, 318–334 (2008).

2. Biteau, B., Hochmuth, C. E. & Jasper, H. JNK activity in somatic stem cells causes loss of tissue homeostasis in the aging Drosophila gut. Cell Stem Cell 3, 442–455 (2008).

3. Li, H., Qi, Y. & Jasper, H. Preventing Age-Related Decline of Gut Compartmentalization Limits Microbiota Dysbiosis and Extends Lifespan. Cell Host Microbe 19, 240–253 (2016).

4. He, D. et al. Gut stem cell aging is driven by mTORC1 via a p38 MAPK-p53 pathway. Nat Commun 11, 37 (2020).

5. Pentinmikko, N. et al. Paneth cell produced Notum attenuates regeneration of aged intestinal epithelium. Nature 571, 398–402 (2019).

6. Guo, L., Karpac, J., Tran, S. L. & Jasper, H. PGRP-SC2 promotes gut immune homeostasis to limit commensal dysbiosis and extend lifespan. Cell 156, 109–122 (2014).

7. Clark, R. I. et al. Distinct Shifts in Microbiota Composition during Drosophila Aging Impair Intestinal Function and Drive Mortality. Cell Rep 12, 1656–1667 (2015).

8. Thevaranjan, N. et al. Age-Associated Microbial Dysbiosis Promotes Intestinal Permeability, Systemic Inflammation, and Macrophage Dysfunction. Cell Host Microbe 21, 455–466.e4 (2017).

9. Katz, D., Hollander, D., Said, H. M. & Dadufalza, V. Aging-associated increase in intestinal permeability to polyethylene glycol 900. Dig Dis Sci 32, 285–288 (1987).

10. Rera, M., Clark, R. I. & Walker, D. W. Intestinal barrier dysfunction links metabolic and inflammatory markers of aging to death in Drosophila. Proc Natl Acad Sci U S A 109, 21528–21533 (2012).

11. Tran, L. & Greenwood-Van Meerveld, B. Age-associated remodeling of the intestinal epithelial barrier. J Gerontol A Biol Sci Med Sci 68, 1045–1056 (2013).

12. Dambroise, E. et al. Two phases of aging separated by the Smurf transition as a public path to death. Sci Rep 6, 23523 (2016).

13. Mitchell, E. L. et al. Reduced Intestinal Motility, Mucosal Barrier Function, and Inflammation in Aged Monkeys. J Nutr Health Aging 21, 354–361 (2017).

14. Buchon, N., Broderick, N. A., Chakrabarti, S. & Lemaitre, B. Invasive and indigenous microbiota impact intestinal stem cell activity through multiple pathways in Drosophila. Genes Dev. 23, 2333–2344 (2009).

15. Smith, P. et al. Regulation of life span by the gut microbiota in the short-lived African turquoise killifish. Elife 6, e27014 (2017).

16. Onuma, T. et al. Recognition of commensal bacterial peptidoglycans defines Drosophila gut homeostasis and lifespan. PLOS Genetics 19, e1010709 (2023).

17. Salazar, A. M. et al. Intestinal Snakeskin Limits Microbial Dysbiosis during Aging and Promotes Longevity. iScience 9, 229–243 (2018).

18. Resnik-Docampo, M. et al. Keeping it tight: The relationship between bacterial dysbiosis, septate junctions, and the intestinal barrier in Drosophila. Fly (Austin) 12, 34–40 (2018).

19. Zhu, Y. et al. Aspirin Positively Contributes to Drosophila Intestinal Homeostasis and Delays Aging through Targeting Imd. Aging Dis 12, 1821–1834 (2021).

20. Rodriguez-Fernandez, I. A., Qi, Y. & Jasper, H. Loss of a proteostatic checkpoint in intestinal stem cells contributes to age-related epithelial dysfunction. Nat Commun 10, 1050 (2019).

21. Wang, L., Ryoo, H. D., Qi, Y. & Jasper, H. PERK Limits Drosophila Lifespan by Promoting Intestinal Stem Cell Proliferation in Response to ER Stress. PLoS Genet 11, e1005220 (2015).

22. Longo, V. D. & Anderson, R. M. Nutrition, longevity and disease: from molecular mechanisms to interventions. Cell 185, 1455–1470 (2022).

23. Wickramasinghe, K., Mathers, J. C., Wopereis, S., Marsman, D. S. & Griffiths, J. C. From lifespan to healthspan: the role of nutrition in healthy ageing. J Nutr Sci 9, e33 (2020).

24. Vinardell, M. P. Age influences on amino acid intestinal transport. Comp Biochem Physiol Comp Physiol 103, 169–171 (1992).

25. Ferraris, R. P., Hsiao, J., Hernandez, R. & Hirayama, B. Site density of mouse intestinal glucose transporters declines with age. Am J Physiol 264, G285–293 (1993).

26. Ferraris, R. P. & Vinnakota, R. R. Regulation of intestinal nutrient transport is impaired in aged mice. J Nutr 123, 502–511 (1993).

27. Tatge, L., Solano Fonseca, R. & Douglas, P. M. A framework for intestinal barrier dysfunction in aging. Nat Aging 1–3 (2023) doi:10.1038/s43587-023-00492-0.

28. Mihaylova, M. M. et al. When a calorie is not just a calorie: Diet quality and timing as mediators of metabolism and healthy aging. Cell Metabolism 35, 1114–1131 (2023).

29. Piper, M. D. et al. A holidic medium for Drosophila melanogaster. Nat Methods 11, 100– 105 (2014).

30. Piper, M. D. W. et al. Matching Dietary Amino Acid Balance to the In Silico-Translated Exome Optimizes Growth and Reproduction without Cost to Lifespan. Cell Metab 25, 1206 (2017).

31. Rera, M. et al. Modulation of longevity and tissue homeostasis by the Drosophila PGC-1 homolog. Cell Metab 14, 623–634 (2011).

32. Babygirija, R. & Lamming, D. W. The regulation of healthspan and lifespan by dietary amino acids. Transl Med Aging 5, 17–30 (2021).

33. Bass, T. M. et al. Optimization of Dietary Restriction Protocols in Drosophila. J Gerontol A Biol Sci Med Sci 62, 1071–1081 (2007).

34. Ja, W. W. et al. Prandiology of Drosophila and the CAFE assay. Proc Natl Acad Sci U S A 104, 8253–8256 (2007).

35. Dutta, D. et al. Regional Cell-Specific Transcriptome Mapping Reveals Regulatory Complexity in the Adult Drosophila Midgut. Cell Rep 12, 346–358 (2015).

36. Krause, S. A., Overend, G., Dow, J. A. T. & Leader, D. P. FlyAtlas 2 in 2022: enhancements to the Drosophila melanogaster expression atlas. Nucleic Acids Res 50, D1010–D1015 (2022).

37. Hu, Y. et al. An integrative approach to ortholog prediction for disease-focused and other functional studies. BMC Bioinformatics 12, 357 (2011).

38. Hu, Y. et al. FlyRNAi.org-the database of the Drosophila RNAi screening center and transgenic RNAi project: 2021 update. Nucleic Acids Res 49, D908–D915 (2021).

39. Goberdhan, D. C. I., Meredith, D., Boyd, C. A. R. & Wilson, C. PAT-related amino acid transporters regulate growth via a novel mechanism that does not require bulk transport of amino acids. Development 132, 2365–2375 (2005).

40. Deng, H., Gerencser, A. A. & Jasper, H. Signal integration by Ca(2+) regulates intestinal stem-cell activity. Nature 528, 212–217 (2015).

41. Colombani, J. et al. A nutrient sensor mechanism controls Drosophila growth. Cell 114, 739–749 (2003).

42. Roman, G., Endo, K., Zong, L. & Davis, R. L. P{Switch}, a system for spatial and temporal control of gene expression in Drosophila melanogaster. Proc Natl Acad Sci U S A 98, 12602–12607 (2001).

43. Osterwalder, T., Yoon, K. S., White, B. H. & Keshishian, H. A conditional tissue-specific transgene expression system using inducible GAL4. Proceedings of the National Academy of Sciences of the United States of America 98, 12596 (2001).

44. Bjordal, M., Arquier, N., Kniazeff, J., Pin, J. P. & Léopold, P. Sensing of amino acids in a dopaminergic circuitry promotes rejection of an incomplete diet in Drosophila. Cell 156, 510–521 (2014).

45. Sun, J. et al. Drosophila FIT is a protein-specific satiety hormone essential for feeding control. Nat Commun 8, 14161 (2017).

46. Yamada, R., Deshpande, S. A., Bruce, K. D., Mak, E. M. & Ja, W. W. Microbes Promote Amino Acid Harvest to Rescue Undernutrition in Drosophila. Cell Rep 10, 865–872 (2015).

47. Keebaugh, E. S., Yamada, R., Obadia, B., Ludington, W. B. & Ja, W. W. Microbial Quantity Impacts Drosophila Nutrition, Development, and Lifespan. iScience 4, 247–259 (2018).

48. Buchon, N., Broderick, N. A., Kuraishi, T. & Lemaitre, B. Drosophila EGFR pathway coordinates stem cell proliferation and gut remodeling following infection. BMC Biol 8, 152 (2010).

49. Zhai, Z., Boquete, J.-P. & Lemaitre, B. Cell-Specific Imd-NF-κB Responses Enable Simultaneous Antibacterial Immunity and Intestinal Epithelial Cell Shedding upon Bacterial Infection. Immunity 48, 897–910.e7 (2018).

50. Fan, X. et al. Rapamycin preserves gut homeostasis during Drosophila aging. Oncotarget 6, 35274–35283 (2015).

51. Schinaman, J. M., Rana, A., Ja, W. W., Clark, R. I. & Walker, D. W. Rapamycin modulates tissue aging and lifespan independently of the gut microbiota in Drosophila. Sci Rep 9, 7824 (2019).

52. Juricic, P. et al. Long-lasting geroprotection from brief rapamycin treatment in early adulthood by persistently increased intestinal autophagy. Nat Aging 2, 824–836 (2022).

53. Lu, J. et al. Sestrin is a key regulator of stem cell function and lifespan in response to dietary amino acids. Nat Aging 1, 60–72 (2021).

54. Na, H.-J. et al. Metformin inhibits age-related centrosome amplification in Drosophila midgut stem cells through AKT/TOR pathway. Mechanisms of Ageing and Development 149, 8–18 (2015).

55. Wu, M., Xiao, H., Shao, F., Tan, B. & Hu, S. Arginine accelerates intestinal health through cytokines and intestinal microbiota. Int Immunopharmacol 81, 106029 (2020).

56. Singh, K. et al. Dietary Arginine Regulates Severity of Experimental Colitis and Affects the Colonic Microbiome. Front Cell Infect Microbiol 9, 66 (2019).

57. Nüse, B. & Mattner, J. L-arginine as a novel target for clinical intervention in inflammatory bowel disease. Explor Immunol. 1, 80–89 (2021).

58. Kim, Y. J. et al. Arginine-mediated gut microbiome remodeling promotes host pulmonary immune defense against nontuberculous mycobacterial infection. Gut Microbes 14, 2073132 (2022).

59. Xiao, W. et al. Glutamate prevents intestinal atrophy via luminal nutrient sensing in a mouse model of total parenteral nutrition. FASEB J 28, 2073–2087 (2014).

60. Hou, Q. et al. Exogenous L-arginine increases intestinal stem cell function through CD90+ stromal cells producing mTORC1-induced Wnt2b. Commun Biol 3, 611 (2020).

61. Ma, S. et al. Cell culture-based profiling across mammals reveals DNA repair and metabolism as determinants of species longevity. eLife 5, e19130 (2016).

62. Edwards, C. et al. Mechanisms of amino acid-mediated lifespan extension in Caenorhabditis elegans. BMC Genet 16, 8 (2015).

63. Tsugawa, Y., Handa, H. & Imai, T. Arginine induces IGF-1 secretion from the endoplasmic reticulum. Biochem Biophys Res Commun 514, 1128–1132 (2019).

64. Alba-Roth, J., Müller, O. A., Schopohl, J. & Werder, K. V. Arginine Stimulates Growth Hormone Secretion by Suppressing Endogenous Somatostatin Secretion. The Journal of Clinical Endocrinology & Metabolism 67, 1186–1189 (1988).

65. Holzenberger, M. et al. IGF-1 receptor regulates lifespan and resistance to oxidative stress in mice. Nature 421, 182–187 (2003).

66. Aguiar-Oliveira, M. H. & Bartke, A. Growth Hormone Deficiency: Health and Longevity. Endocrine Reviews 40, 575–601 (2019).

67. Canfield, C.-A. & Bradshaw, P. C. Amino acids in the regulation of aging and aging-related diseases. Translational Medicine of Aging 3, 70–89 (2019).

68. Carroll, B. et al. Control of TSC2-Rheb signaling axis by arginine regulates mTORC1 activity. Elife 5, e11058 (2016).

69. Hoffman, J. M. et al. Effects of age, sex, and genotype on high-sensitivity metabolomic profiles in the fruit fly, Drosophila melanogaster. Aging Cell 13, 596–604 (2014).

70. Bayliak, M. M. et al. Dietary L-arginine accelerates pupation and promotes high protein levels but induces oxidative stress and reduces fecundity and life span in Drosophila melanogaster. J Comp Physiol B 188, 37–55 (2018).

71. Govindaraju, T., Sahle, B. W., McCaffrey, T. A., McNeil, J. J. & Owen, A. J. Dietary Patterns and Quality of Life in Older Adults: A Systematic Review. Nutrients 10, 971 (2018).

72. Volpi, E., Mittendorfer, B., Rasmussen, B. B. & Wolfe, R. R. The response of muscle protein anabolism to combined hyperaminoacidemia and glucose-induced hyperinsulinemia is impaired in the elderly. J Clin Endocrinol Metab 85, 4481–4490 (2000).

73. Katsanos, C. S., Kobayashi, H., Sheffield-Moore, M., Aarsland, A. & Wolfe, R. R. Aging is associated with diminished accretion of muscle proteins after the ingestion of a small bolus of essential amino acids2. The American Journal of Clinical Nutrition 82, 1065– 1073 (2005).

74. Katsanos, C. S., Kobayashi, H., Sheffield-Moore, M., Aarsland, A. & Wolfe, R. R. A high proportion of leucine is required for optimal stimulation of the rate of muscle protein synthesis by essential amino acids in the elderly. Am J Physiol Endocrinol Metab 291, E381–387 (2006).

75. Kabil, H., Kabil, O., Banerjee, R., Harshman, L. G. & Pletcher, S. D. Increased transsulfuration mediates longevity and dietary restriction in Drosophila. Proc Natl Acad Sci U S A 108, 16831–16836 (2011).

76. Lewis, E. A new standard food medium. Drosophila Information Service 117–118 (1960).

77. Bakula, M. The persistence of a microbial flora during postembryogenesis of Drosophila melanogaster. Journal of Invertebrate Pathology 14, 365–374 (1969).

78. Claesson, M. J. et al. Composition, variability, and temporal stability of the intestinal microbiota of the elderly. Proc Natl Acad Sci U S A 108 Suppl 1, 4586–4591 (2011).

