## Supplemental defined media protocol for "Drosophila Undigested Metabolite Profiling reveals age related loss of intestinal amino acid transport regulates longevity"

Amounts of amino acids taken from the exome matched diet (Piper, 2017). Total mass of amino acids in 1 L = 21.4g for the standard diet. We also use the addition of 50% more vitamin solution, which in the original paper (Piper, 2014) resulted in improved proportions of flies surviving from egg to adult. Quantities given below make 1 litre of food. Quantities are given for both the standard media and Aged Amino Acid media. Reagents were sourced as described in the original publications and are available online from DOI: [10.1038/protex.2013.082](https://doi.org/10.1038/protex.2013.082) table 1 (reorderlist.pdf in supplemental)

### *Quick guide*

- To cook, combine ingredients from buffer 1, and heat to a boil in the microwave.
- Allow to cool to 65ºC.
- Whilst cooling, measure out the other ingredients (buffer 2, excluding nipagin and propionic acid) in the laminar flow hood (to ensure the stock solutions remain sterile)
- Once buffer 1 is cooled to 65ºC, add buffer 2, the nipagin and propionic acid, and mix thoroughly.

#### Buffer 1:

| | **Standard** | **Aged Amino Acid** |
|---|---|---|
| ***Agar*** | 20 (g) | 20 (g) |
| ***Sucrose*** | 17.12 (g) | 17.12 (g) |
| ***Isoleucine*** | 1.12 (g) | 1.81 (g) |
| ***Leucine*** | 2.03 (g) | 2.65 (g) |
| ***Tyrosine*** | 0.93 (g) | 2.06 (g) |
| ***Acetate buffer*** | 100 (mL) | 100 (mL) |
| ***CaCl2*** | 1 (mL) | 1(mL) |
| ***CuSO4*** | 1 (mL) | 1 (mL) |
| ***FeSO4*** | 1 (mL) | 1 (mL) |
| ***M(g)SO4*** | 1 (mL) | 1 (mL) |
| ***MnCl2*** | 1 (mL) | 1 (mL) |
| ***ZnSO4*** | 1 (mL) | 1 (mL) |
| ***Cholesterol*** | 15 (mL) | 15 (mL) |
| ***Milli Q water*** | 685 (mL) | 624.5 (mL) |

#### Buffer 2:

| | **Standard** | **Aged Amino Acid** |
|---|---|---|
| ***Amino Acids*** | 121 (mL) | 181.6 (mL) |
| ***Cysteine*** | 6.8 (mL) | 6.8 (mL) |
| ***NaGlu*** | 15.1 (mL) | 15.1 (mL) |
| ***Sodium Folate*** | 1 (mL) | 1 (mL) |
| ***Vitamins*** | 21 (mL) | 21 (mL) |
| ***Other nutrients*** | 8 (mL) | 8 (mL) |
| ***Propionic Acid*** | *6 (mL)* | *6 (mL)* |
| ***10% Nipagin*** | *15 (mL)* | *15 (mL)* |

### *Detailed instructions*

#### *Preparation*

- Remove cholesterol from fridge and heat to solubilise – we sit it in hot water and agitate occasionally.
- Remove iron solution from freezer to thaw. **Do not allow the FeSO4 to sit at room temperature too long or it will oxidise.**

#### *Cooking buffer 1*

- Measure the sucrose (Sigma) into a beaker.
- Use the fine balance and weigh the Leucine, Isoleucine and Tyrosine into a weighing boat. Transfer into beaker.
- Add the MilliQ water (rinse the weighing boat)
- Add the cholesterol and metal ion solutions and acetate buffer, then mix. The solution will turn cloudy upon addition of cholesterol
- Microwave until the agar is melted and the solution boils

Allow to cool to 65°C (ideally on a stirrer). *We incubate in a water bath set to 50°C as this gives us time to weigh out buffer 2 components without allowing buffer 1 to set.*

- **In the laminar flow hood**, measure out all the components of buffer 2, excluding the nipagin and propionic acid, into a separate beaker
    - This is to ensure the solutions remain sterile.
- Once buffer 1 has cooled to 65°C, add buffer 2 and the nipagin and propionic acid and mix thoroughly.
- Dispense into vials

---

### Sterile Defined food

- Measure all of buffer 1* into a duran of greater volume than the media.
- Add additional MilliQ water to the same volume as the Nipagin and Propionic Acid, as these are omitted from sterile food
- Add a magnetic flea to the bottle, and autoclave at 121°C for 15 minutes.
    - Ensure the lid is not tightly closed.
    - Also autoclave vials and stoppers
- Cool on stirrer in laminar flow (thermometer sterilised in 80% EtOH)
- Measure out buffer 2 in laminar flow (excluding nipagin and propionic acid) into a sterile container.
- Once buffer 1 has cooled to 65°C, add buffer 2 and allow to mix in thoroughly.
- Dispense under laminar flow (or other suitably sterile conditions into sterile vials. Stopper with sterile plugs.
- **Do not** add nipagin or acid mix to the autoclaved food (as they are not sterile)
- Ensure that the food is fully set prior to use
- *if the food does not set, autoclave buffer 1 without the acetate buffer. Add acetate buffer with buffer 2 (filter sterilise or autoclave the acetate buffer separately before adding it).

### Stock solutions

#### Amino Acid solution:

To make 1L of amino acid solution, combine the following amounts of each amino acid. Quantities are given for the standard diet and the Aged Amino Acid diet.

| *Essential Amino Acids* | *Standard* | *Aged Amino Acid* | |
|---|---|---|---|
| *Phenylalanine* | 8.32 | 10.91 | g |
| *Histidine* | 5.4 | 9.11 | g |
| *Lysine* | 11.26 | 8.85 | g |
| *Methionine* | 4.98 | 5.0 | g |
| *Arginine* | 13.46 | 14.07 | g |
| *Threonine* | 9.13 | 8.45 | g |
| *Valine* | 9.9 | 11.99 | g |
| *Tryptophan* | 2.65 | 9.15 | g |

| *Non-Essential Amino Acids* | Standard | Aged Amino Acid | |
|---|---|---|---|
| *Alanine* | 9.09 | 9.24 | g |
| *Aspartic Acid* | 9.67 | 7.39 | g |
| *Glycine* | 6.33 | 6.44 | g |
| *Asparagine* | 8.50 | 10.41 | g |
| *Proline* | 8.07 | 9.49 | g |
| *Glutamine* | 9.25 | 14.03 | g |
| *Serine* | 11.37 | 11.15 | g |

Add 800 mL mQ water protect from light and leave on stirrer until dissolved. Once fully dissolved, check final volume, add more mQ water if necessary to make 1L solution.
Use 500 mL BD plastipak syringe and 0.4uM filter to sterilise and dispense the solution into a sterile (autoclaved) Duran bottle in a laminar flow hood.
Store at 4°C

#### *Cysteine*

| **Cysteine** | **Desired Volume (ml):** | **40** |
|---|---|---|
| **Cysteine** | 2 | g |

Make up to volume with MilliQ water. It **dissolves almost immediately**.
Once fully dissolved, use 500 mL BD plastipak syringe and 0.4uM filter to sterilise and dispense the solution into a sterile (autoclaved) Duran bottle in a laminar flow hood.
Store at 4°C

#### *Sodium Glutamate (NaGlu)*

| **NaGlu** | **Desired Volume (ml):** | **100** |
|---|---|---|
| **Glutamic Acid: Standard** | 10 | g |
| **Glutamic Acid: Aged** | 32.8 | g |

Make up to volume with MilliQ water. It **dissolves almost immediately**.
Once fully dissolved, use 500 mL BD plastipak syringe and 0.4uM filter to sterilise and dispense the solution into a sterile (autoclaved) Duran bottle in a laminar flow hood.
Store at 4°C

### Metal Ion solutions:

Weigh out and dissolve in MilliQ water. For some solutions this may require leaving on a stirrer. Once fully dissolved aliquot into 1.5mL microcentrifuge tubes and store at -20°C.
As some of these solutions must be made in large volumes to prevent such large weighing error, they can be made aliquoted into 50mL falcon tubes and stored at -20°C to be made into 1.5mL aliquots as required.
Alternatively, filter sterilise solutions (excluding $FeSO_4$) and store at 4°C.

| **MnCl2** | **Desired Volume (ml):** | **500** |
|---|---|---|
| **Manganese Chloride Tetrahydrate** | 0.5 | g |
| **Make up to volume with MilliQ Water** | | |

| **ZnSO4** | **Desired Volume (ml):** | **100** |
|---|---|---|
| **Zinc Sulfate Heptahydrate** | 2.5 | g |
| **Make up to volume with MilliQ Water** | | |

| **FeSO4** | **Desired Volume (ml):** | **100** |
|---|---|---|
| **Iron(II) Sulfate Heptahydrate** | 2.5 | g |
| **Make up to volume with MilliQ Water** | | |

| **CuSO4** | **Desired Volume (ml):** | **200** |
|---|---|---|
| **Copper(II) Sulfate Heptahydrate** | 0.5 | g |
| **Make up to volume with MilliQ Water** | | |

| **MgSO4** | **Desired Volume (ml):** | **50** |
|---|---|---|
| **Magnesium Sulfate** | 12.5 | g |
| **Make up to volume with MilliQ Water** | | |

| **CaCl2** | **Desired Volume (ml):** | **50** |
|---|---|---|
| **Calcium Chloride** | 12.5 | g |
| **Make up to volume with MilliQ Water** | | |

### Vitamins and other solutions

#### *Folic Acid*

| Folic Acid | Desired Volume (ml): | 500 |
|---|---|---|
| **Folic Acid** | 0.25 | g |

Make up to volume with MilliQ water. If required, bring into solution by drop-wise addition of 2 N NaOH solution.

*N = normality. For NaOH, this is the same as molarity because its acidity is 1. Therefore 2N = 2M*

Store at -20°C long-term, once thawed store at 4°C

#### *Cholesterol*

Cholesterol powder is stored at -20C

| Cholesterol | Desired Volume (ml): | 200 |
|---|---|---|
| **Cholesterol** | 4 | g |
| **Make up to volume with 100% EtOH** | | |

Dissolve in ethanol, aliquot into 15mL falcon tube, and store at 4°C. Rinse beaker with ethanol
Before use, incubate on shaker at 37°C for 30-60 minutes to re-dissolve

#### *Acetate Buffer*

| Acetate Buffer | Desired Volume (ml): | 1000 |
|---|---|---|
| **Acetic Acid** | 30 | ml |
| **Potassium Phosphate Monobasic** | 30 | g |
| **Sodium Bicarbonate** | 10 | g |

Weigh the dry ingredients into a 1L sterile duran bottle.
**Slowly** add the 600 mL of water. This will effervesce, so be careful not to overflow.
Add the acetic acid and then top up to 1000mL
Adjust the pH to 4
Store at 4°C

#### *Nucleic acids and lipid related metabolites*

| Nucleic acids and lipid-related metabolites | Desired Volume (ml): | 200 |
|---|---|---|
| **Choline Chloride** | 1.25 | g |
| **Myo-Inositol** | 0.126 | g |
| **Inosine** | 1.626 | g |
| **Uridine** | 1.5 | g |

Make up to volume with MilliQ water, protect from light and leave on stirrer until dissolved.
Once fully dissolved, use 500 mL BD plastipak syringe and 0.4uM filter to sterilise and dispense the solution into a sterile (autoclaved) Duran bottle in the laminar flow hood.
Store at 4°C

*Vitamin solution*

| Vitamin Solution | Desired Volume (ml): | 1000 |
|---|---|---|
| **Thiamine** | 100 | mg |
| **Riboflavin** | 50 | mg |
| **Nicotinic Acid** | 600 | mg |
| **Ca Pantothenate** | 755 | mg |
| **Pyridoxine** | 125 | mg |
| **Biotin** | 10 | mg |

Make up to volume with MilliQ water, protect from light and leave on stirrer until dissolved.
Once fully dissolved, use syringe and 0.4uM filter to sterilise and dispense the solution into a sterile (autoclaved) Duran, or 50mL falcon tubes, in the laminar flow hood.
Store at -20°C long-term, once thawed store at 4°C
